# Beta cell reactivity defines disease-relevant pancreatic CD8 T cells in type 1 diabetes

**DOI:** 10.64898/2026.06.08.729940

**Authors:** Amanda M. Anderson, Laurie G. Landry, Jessie M. Barra, Kristen L. Wells, Ali H. Shilleh, Roberto Castro-Gutierrez, Andrew M. Ladd, Megan DeNicola, Jenny Aurielle B. Babon, Roberto Mallone, Sally C. Kent, Aaron W. Michels, Holger A. Russ, Maki Nakayama

**Affiliations:** Barbara Davis Center for Childhood Diabetes, University of Colorado School of Medicine, Aurora, CO, USA; University of Florida Diabetes Institute, University of Florida, Gainesville, FL, USA; Department of Pharmacology & Therapeutics, College of Medicine, University of Florida, Gainesville, FL, USA; Diabetes Center of Excellence, Department of Medicine, University of Massachusetts Chan Medical School, Worcester, MA, USA; Université Paris Cité, Institut Cochin, CNRS, Inserm, Paris, France; Assistance Publique Hôpitaux de Paris, Cochin Hospital, Department of Diabetology and Clinical Immunology, Paris, France; Indiana Biosciences Research Institute, Indianapolis, IN, USA; Department of Pediatrics, University of Colorado School of Medicine, Aurora, CO, USA; Department of Immunology and Microbiology, University of Colorado School of Medicine, Aurora, CO, USA

## Abstract

Type 1 diabetes (T1D) is characterized by immune-mediated destruction of pancreatic beta cells, yet the properties that distinguish disease-associated CD8 T cells from other pancreatic resident T cells remain incompletely defined. In this study, we analyzed CD8 T cell receptor (TCR) clonotypes isolated from the pancreas of organ donors with and without T1D and assessed their reactivity to beta cells using stem cell-derived beta-like cells. We found that highly beta cell-reactive CD8 T cells were selectively present in the pancreas of T1D donors but were largely absent from donors without T1D. In contrast, virus-specific CD8 T cells were detected in pancreata of donors with and without T1D and showed no evidence of cross-reactivity to beta-like cells, indicating that pancreatic residency alone does not confer beta cell specificity. Among beta cell-reactive CD8 T cells in T1D, reactivity to native peptides from major islet proteins other than preproinsulin was rare. Thus, despite beta cell specificity as a hallmark of T1D, T cells reactive to native islet proteins other than preproinsulin do not infiltrate the islets. These results identify beta cell reactivity as a key functional feature separating T1D-associated CD8 T cells from other pancreatic T cells. This functional definition of pathogenic T cells offers a framework for understanding selective beta cell loss and for developing approaches to monitor and therapeutically target disease-relevant CD8 T cells.

## Introduction

Insulin-secreting beta cells within the pancreatic islets are destroyed by immune-mediated mechanisms in which T cells play a central role during the development of type 1 diabetes (T1D) (*1–4*). Pancreatic islets contain multiple endocrine cell types; however, evidence of autoimmune attack is largely restricted to beta cells (*5, 6*). It remains unclear why beta cells, among the various endocrine cell types, are selectively attacked by the adaptive immune system. Antigens recognized by T cells may represent a key determinant of this tissue specificity, and therefore, extensive efforts have been made to identify the antigenic targets of T cells involved in the pathogenesis of T1D (*7–9*).

Both CD4 and CD8 T cells play crucial roles in T1D development. Strong genetic associations within the HLA locus, particularly HLA class II genes, highlight the essential role of CD4 T cells (*10, 11*). Likewise, HLA class I (HLA-I) genes are also linked to T1D risk, and notably, the rate of progression to clinical T1D correlates with specific HLA-I alleles, implicating CD8 T cells in beta cell destruction through recognition of epitopes presented by risk-associated HLA-I molecules (*10–12*). Indeed, we and others have demonstrated the dominant presence of CD8 T cells within the pancreas of T1D organ donors, with markedly higher frequencies in individuals with clinical T1D compared to islet autoantibody-positive or non-diabetic donors (*13–17*). Histological analyses further reveal elevated HLA-I expression on beta cells in the pancreata of T1D organ donors, especially involving HLA-B molecules (*13, 18, 19*). Importantly, a higher abundance of circulating exhausted CD8 T cells correlates with slower disease progression (*20*). Consistent with this, stage 2 individuals treated with anti-CD3 monoclonal antibody, which delays the time to clinical stage 3 T1D onset, exhibit an increased proportion of CD8 T cells expressing exhaustion markers compared with placebo-treated arms (*21*). Collectively, these findings support a critical role of CD8 T cells in T1D pathogenesis and disease progression.

Given the central involvement of CD8 T cells in beta cell destruction, considerable effort has been devoted to defining the antigen specificity of CD8 T cells in the pancreas. Identifying epitopes recognized by these cells provides key insights into the mechanisms by which self-tolerance is broken and beta cell autoimmunity evolves over time, thereby guiding the design and appropriate timing of targeted immunotherapies. Historically, epitopes recognized by CD8 T cells were first identified from peripheral blood of individuals with and without T1D (*7, 22–24*). Subsequently, CD8 T cells specific for the same epitopes, such as preproinsulin (PPI) 15-24 and zinc-transporter 8 (ZnT8) 186-194, were confirmed to be present in the islets of T1D organ donors using peptide-HLA multimer-based histological analyses (*13, 15*). Later, CD8 T cell lines were established from the pancreatic islets of a T1D donor and were tested for their specificity for several islet peptides using multimers; however, most cells in these lines were not stained by these peptide-loaded multimers (*14*). Thereafter, we generated transductant cells expressing T cell receptors (TCRs) identified from pancreatic and islet T cells and comprehensively tested their responses to native PPI-derived peptides presented by autologous HLA-I molecules (*17*). We found that T1D donors harbor CD8 T cells reactive to native PPI peptides presented by various HLA-I molecules in the pancreas, whereas among TCRs identified from non-diabetic donors, only one showed very weak reactivity to a PPI peptide. A recent study focusing on an HLA-C allele further identified epitopes within the insulin A-chain recognized by CD8 T cells in the islets of T1D donors (*25*). However, despite decades of study, numerous pancreatic CD8 T cells remain of unknown antigen specificity, and importantly, their reactivity to beta cells has not been fully defined. Thus, the antigenic landscape of pancreatic CD8 T cells involved in T1D pathogenesis remains incompletely resolved.

What distinguishes autoreactive T cells that successfully enter and respond within the human pancreas—and how this is shaped by antigen specificity—remains a central question in T1D pathogenesis. A key unresolved question is whether progression to clinical disease reflects a broad loss of tolerance across the islet proteome or the selective enrichment of T cell clones with defined antigen recognition properties. In this study, we sought to define the antigenic rules governing pancreatic CD8 T cell autoreactivity in human T1D. By systematically examining hundreds of pancreatic CD8 TCRs from donors with and without T1D, we found that, among native islet peptides tested, autoreactivity is not broadly distributed across major islet proteins but is instead preferentially directed toward PPI. Furthermore, highly beta cell-reactive T cells were observed exclusively in T1D donors and were not detected in non-diabetic donors despite the presence of pancreatic T cells, indicating that entry of islet-reactive T cells into the pancreas is tightly restricted under non-diabetic conditions. Thus, beta cell reactivity is not a general feature of pancreatic resident T cells, but a disease-associated property selectively observed in T1D.

Here we demonstrate the selective accumulation of beta cell–reactive CD8 T cells as a hallmark of T1D, representing a disease-specific component of the pancreatic immune repertoire, with reactivity to native peptides preferentially directed toward PPI, alongside a population of beta cell-reactive T cells not accounted for by native peptide recognition.

## Results

### Single anti-GAD antibody positivity is insufficient to confer anti-insulin autoimmunity by CD8 T cells

This study analyzed 5,651 CD8 T cells isolated from the islets or pancreata of organ donors with T1D (n=16), single anti-GAD antibody positive (n=4), and non-diabetic controls (n=10), for which in-frame TCR alpha and/or beta chain sequences were successfully identified (**Table 1**, **Supplementary Table 1**). Among these clonotypes, 542 unique TCRs derived from 25 donors (13 T1D, 4 single anti-GAD antibody positive, and 8 non-diabetic control donors), most of which were identified from two or more cells, were expressed in T-hybridoma cells lacking endogenous TCR expression to assess antigen specificity (**Supplementary Table 1**). Although a subset of these TCRs had been evaluated previously for reactivity to peptides derived from PPI and insulin-DRiP (defective ribosomal product) (*17*), all newly generated TCR transductants were screened for specificity to PPI and insulin-DRiP using the same experimental strategy, thereby enabling direct comparison of antigen-reactive CD8 T cell frequencies across donor groups and reactivity to all other antigens examined in the present work.

**Table 1.** Summary of donors included in the TCR and antigen analyses.

|  | T1D (n=16) | AAb (n=4) | non-T1D (n=10) |
| --- | --- | --- | --- |
| Age, years (Range (Median)) | 3 to 28 (18.5) | 19 to 29 (20.5) | 13 to 34 (25.5) |
| Diabetes duration, years (Range (Median)) | 0 to 15 (2) | - | - |
| Gender: Female % (number) | 56% (9/16) | 50% (2/4) | 30% (3/10) |
| Islet autoantibodies |  |  |  |
| Insulin % (number) | 67% (10/15) | 0% (0/4) | 0% (0/4) |
| GAD65 % (number) | 47% (7/15) | 100% (4/4) | 0% (0/4) |
| IA-2 % (number) | 47% (7/15) | 0% (0/4) | 0% (0/4) |
| ZnT8 % (number) | 33% (5/15) | 0% (0/4) | 0% (0/4) |
| Number of T cells with TCR sequences identified (Range (Median)) | 46 to 403 (202) | 54 to 384 (217.5) | 62 to 242 (139) |
| Unique in-frame alpha identified (Range (Median)) | 21 to 278 (115) | 41 to 277 (102.5) | 41 to 125 (63) |
| Unique in-frame beta identified (Range (Median)) | 16 to 232 (110) | 36 to 262 (94.5) | 43 to 123 (64) |
| Frequency of T cells responded to antigens (Range (number of individuals with responses)) |  |  |  |
| % PPI* | 0% to 72% (12/13) | 0% | 0% to 2% (2/8) |
| % insulin-DRiP* | 0% to 18% (3/13) | 0% | 0% |
| % GAD65 | 0% | not tested | not tested |
| % IAPP | 0% | not tested | not tested |
| % CHGA | 0% to 2% (1/13) | not tested | not tested |
| % ZNT8 | 0% | not tested | not tested |
| % IGRP | 0% to 3% (1/13) | not tested | not tested |
| % Influenza A | 0% to 2% (3/13) | 0% to 1% (1/4) | 0% to 31% (4/8) |
| % EBV | 0% to 2% (2/13) | 0% | 0% to 3% (1/8) |
| % CMV | 0% | 0% | 0% to 1% (1/8) |
| % Stem cell-derived beta cells | 0% to 76% (7/9) | 0% | 0% to 14% (1/6) |

\* including samples analyzed previously (17)

In the dataset obtained from the newly generated TCR transductants, we confirmed results consistent with prior observations regarding reactivity to PPI and insulin-DRiP in T1D and non-diabetic control donors: 42 and 2 out of 168 TCRs derived from T1D donors were reactive to PPI and insulin-DRiP, respectively (**Supplementary Figure 1, Supplementary Tables 2 and 3**). In contrast, only one TCR clonotype among 63 TCRs detected from pancreas tissues of non-diabetic control donors recognized PPI (**Supplementary Figure 1, Supplementary Tables 2 and 3**). Notably, none of the 70 TCRs from four single anti-GAD antibody-positive donors responded to either PPI or insulin-DRiP peptides (**Supplementary Table 2**). The frequencies of CD8 T cells expressing PPI- and insulin-DRiP-reactive TCR clonotypes in the pancreata of individual donors without overt diabetes (i.e., non-diabetic control and anti-GAD antibody-positive donors) were significantly lower than those in T1D donors (**Figure 1A**, **Table 1**). These results demonstrate that PPI- and insulin-DRiP-reactive CD8 T cells are a hallmark of the pancreas in T1D donors and that the presence of single anti-GAD antibody may be insufficient to confer PPI and insulin-DRiP-reactive CD8 T cell reactivity in the pancreas.

**Figure 1.**
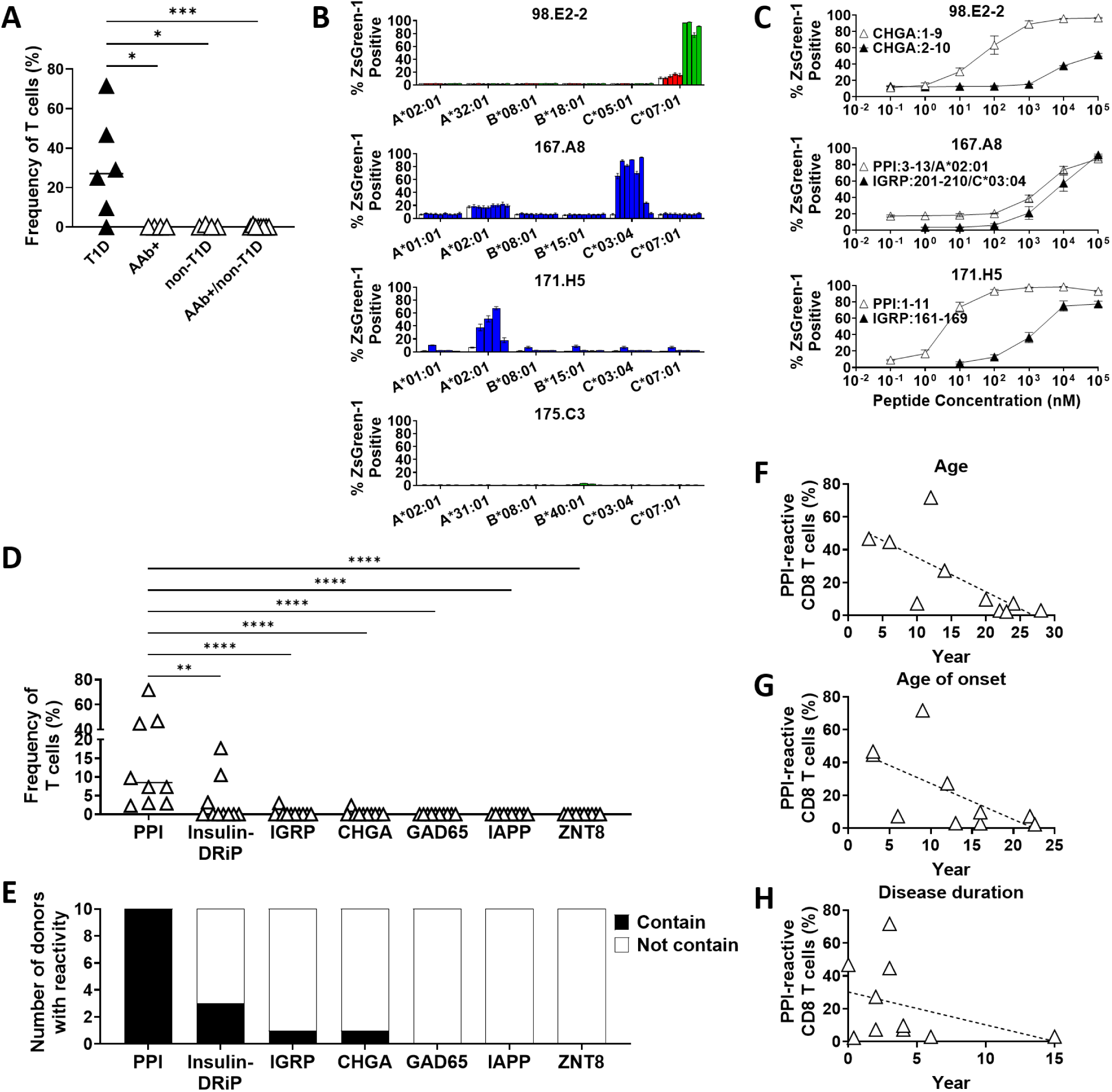
CD8 T cell responses to native islet peptides. (A) Frequencies of CD8 T cells expressing TCRs reactive to PPI peptides are shown for individual donors. Each point represents the proportion of PPI-reactive T cells among all TCR-expressing T cells tested for PPI reactivity in that donor. Frequencies across donor groups (T1D, autoantibody-positive, and non-diabetic controls) were compared using the Mann–Whitney test. *p < 0.05, ***p < 0.005. (B) TCR transductants that responded to truncated peptide pools in the initial screen were tested with individual peptides from the corresponding pools using K562 cells expressing single donor-matched HLA-I molecules to confirm peptide reactivity and define HLA restriction. Responses are shown as the percentage of ZsGreen1-positive cells following peptide stimulation. (C) Dose-response analyses of TCR transductants that responded to defined peptides in the indicated HLA class I context (as identified in panel B). TCR transductants were stimulated with cognate peptides across a range of concentrations in the presence of K562 cells expressing the indicated HLA-I molecules (C*07:01 for 98.E2-2; A*02:01 or C*03:04 for 167.A8; A*02:01 for 171.H5). Activation was assessed by ZsGreen1 expression driven by an NFAT reporter, and mean values ± SEM from two or three independent experiments are shown. (D) Frequencies of CD8 T cells expressing TCRs reactive to native islet peptides are shown for individual T1D donors. Each point represents the proportion of reactive T cells among all T cells expressing tested TCRs in that donor. Frequencies across antigen specificities were compared using the Mann–Whitney test. **p < 0.01, ****p < 0.0001. (E) Number of T1D donors (n = 10) with detectable CD8 T cell responses to native islet peptides. Bars indicate the number of donors with (black) or without (white) detectable responses for each antigen specificity. (F)-(H) Frequencies of PPI-reactive CD8 T cells were plotted against age (F), age of onset (G), and disease duration (H). Statistical significance was assessed using two-tailed Spearman correlation. Age: r = -0.7356, P = 0.0188; age of onset: r = -0.7339, P = 0.0191; disease duration: r = -0.3129, P = 0.3739. Dashed lines indicate linear regression fits shown for visualization.

### Reactivity to native peptides of major islet proteins is rarely detected

We next evaluated the reactivity of TCRs to native peptides derived from major islet proteins including glutamic acid decarboxylase-65 (GAD65), islet amyloid polypeptide (IAPP), chromogranin A (ChgA), ZnT8, and glucose-6-phosphatase 2 (IGRP). We tested 267 TCR clonotypes identified from 10 T1D donors for responses to 1,820 truncated peptide pools, each consisting of 8- to 11-mer peptides ending at the same position within the respective protein sequence, in the presence of antigen-presenting cells (APCs) expressing autologous HLA-A, -B, and -C molecules. Thus, this assay design enabled a comprehensive assessment of TCR specificity to native peptides derived from these five proteins presented by any HLA-I molecules expressed by individual donors. Despite this extensive screening, only a few TCRs responded to any of the truncated peptide pools. The initial screening detected four TCRs that responded to one or more of the 1,820 truncated peptide pools (**Supplementary Table 4**,). Further confirmation assays using newly synthesized, high-purity peptides in the presence of APCs expressing individual HLA-I molecules verified HLA-specific responses by only three TCRs, one responding to ChgA and two responding to IGRP (**Figure 1B**). Interestingly, the two IGRP-reactive TCRs were also cross-reactive with PPI peptides, one of which (TCR 167.A8) responded to each cognate peptide presented by different HLA molecules (**Figure 1C, Supplementary Tables 3 and 4**), thus exhibiting a promiscuous antigen recognition pattern.

### CD8 T cells reactive to native peptides derived from major islet proteins other than preproinsulin are rare in the pancreas of T1D organ donors

Overall, when combined with previously published data analyzing reactivity to PPI and insulin-DRiP (*17*), analysis of ten T1D donors for CD8 T cell reactivity to PPI, insulin-DRiP, GAD65, IAPP, ChgA, ZnT8, and IGRP revealed that all ten donors had at least one TCR clonotype reactive to PPI. These PPI-reactive clonotypes were expressed by 2 – 72% of CD8 T cells analyzed in the pancreas of each donor, and insulin-DRiP-reactive TCRs were detected in three of the 10 donors, accounting for 3 – 18% of CD8 T cells in the pancreas (**Figure 1D and 1E**, **Table 1, Supplementary Table 4**). Notably, the frequency of PPI-reactive T cells was inversely correlated with age and age of onset, but not with disease duration (**Figure F-H**). In contrast, only a small proportion of CD8 T cells reactive to ChgA or IGRP were detected in individual donors, and none of the 267 TCR clonotypes analyzed reacted to native peptides derived from GAD65, ZnT8, and IAPP (**Figure 1D and 1E**, **Table 1, Supplementary Table 4**). Thus, in contrast to the frequent detection of PPI- and insulin-DRiP-reactive CD8 T cells, reactivity to other native islet proteins was detected only sporadically. These results indicate that CD8 T cell entry into the pancreas driven by recognition of other native islet proteins studied, namely GAD65, IAPP, ChgA, ZnT8, and IGRP, is largely limited within individuals having clinical T1D.

### Pancreatic CD8 TCR repertoires contain clonotypes identical to those in the public TCR database

Screening of the 267 TCRs expressed by 932 CD8 T cells in the pancreas of ten T1D donors identified antigen specificity for 28% of T cells, most of which were directed to PPI; however, the targets of nearly three quarters of CD8 T cells remained unknown. To predict the specificity of these orphan TCR clonotypes, we searched for TCRs identical to those listed in a public TCR database, the Immune Epitope Database & Tools (IEDB, https://www.iedb.org/). Among 3,174 alpha and 3,084 beta unique TCR clonotypes identified from CD8 T cells in the pancreas of donors with and without T1D (**Supplementary Table 1**), 13 alpha and 13 beta clonotypes were identical to IEDB public TCRs supported by two or more publications (**Supplementary Table 5**). All IEDB public TCRs shared with those in the pancreas were reported to recognize viral peptides, predominantly TCRs recognizing an influenza A virus matrix protein peptide, FluMP:58-66, presented by HLA-A2 (**Figure 2A**).

**Figure 2.**
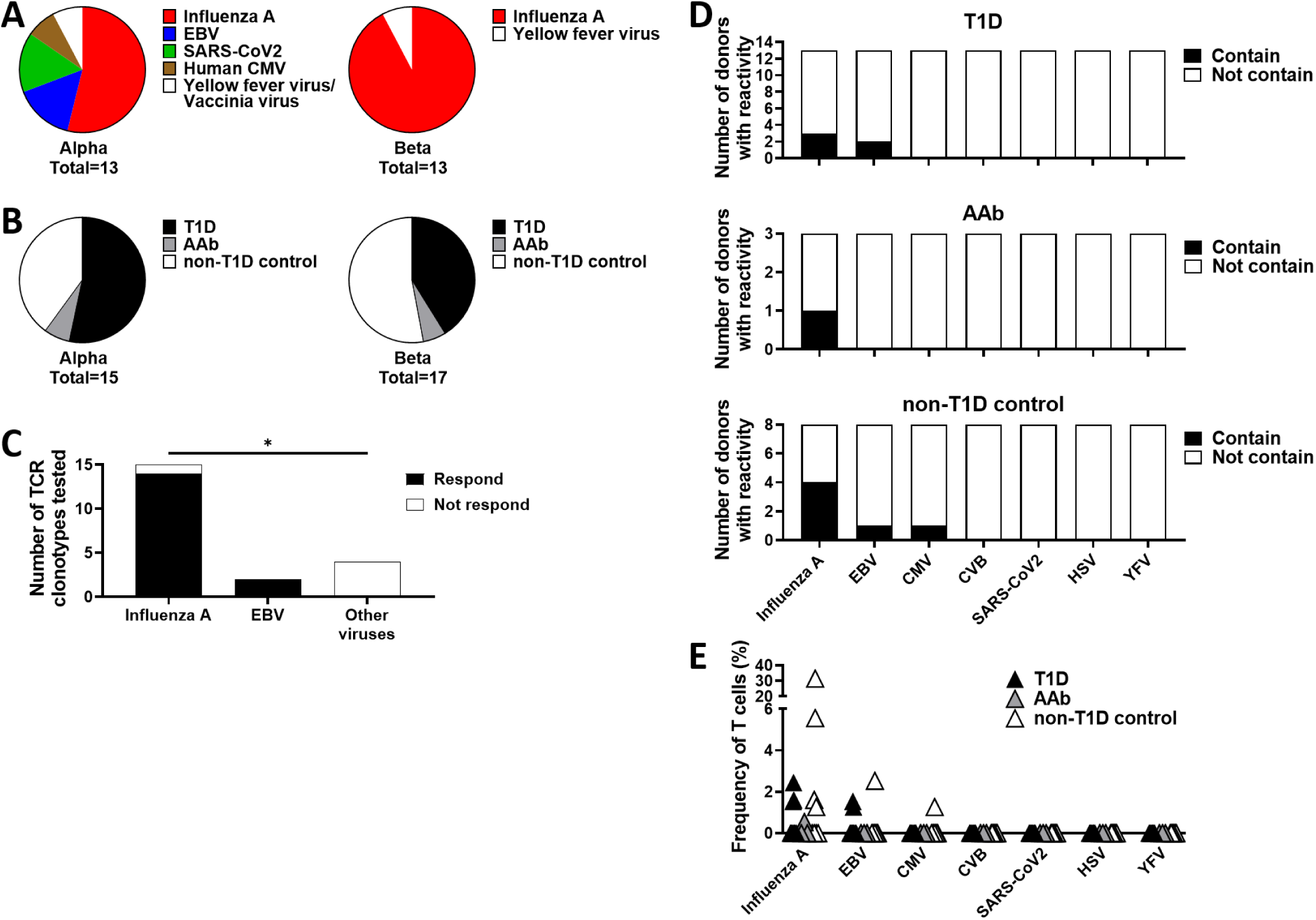
CD8 T cell responses to viral peptides. (A) TCR clonotypes identified from the pancreas were compared to publicly available TCR sequences in the Immune Epitope Database (IEDB). A total of 13 alpha chain and 13 beta chain clonotypes showed exact matches based on full V(D)J sequence identity within each chain. The distribution of source organisms for epitopes associated with the matched TCRs is shown. (B) Donor origin of TCR clonotypes shared with the public database is shown. Among shared clonotypes (15 alpha chains, including 2 detected in multiple donors, and 17 beta chains, including 3 detected in multiple donors; see Supplementary Table 5), the proportions derived from T1D, autoantibody-positive (AAb), and non-diabetic control donors are indicated. (C) Full TCR clonotypes (paired alpha-beta sequences identified from the pancreas) sharing either alpha or beta chains with public TCRs were expressed in T-hybridoma cells and tested for reactivity to cognate peptides reported in the IEDB. The number of TCRs responding to peptides derived from influenza A, EBV, or other viruses is shown. Frequencies of responding TCR clonotypes were compared between influenza A and other viral categories using a two-sided Fisher’s exact test (*P = 0.013). (D) Number of donors with detectable CD8 T cell responses to viral peptides is shown for each donor group (T1D, autoantibody-positive, and non-diabetic controls) across the indicated viral specificities. Bars indicate donors with or without detectable responses. (E) Frequencies of CD8 T cells expressing viral peptide-reactive TCRs are shown for individual donors, grouped by viral specificity. Each point represents the proportion of reactive T cells among all T cells expressing tested TCRs in that donor (black triangles: T1D; gray triangles: autoantibody-positive; white triangles: non-diabetic controls).

These shared TCR clonotypes were present in the pancreas of both T1D and non-diabetic donors (**Figure 2B, Supplementary Table 5**), and all except one clonotype were found in donors carrying the same HLA-I allele that restricts the specificity of the corresponding public TCRs in IEDB (**Supplementary Table 5**). Because these shared clonotypes are not necessarily paired with identical alpha or beta chains between public and pancreatic TCRs, we generated T-hybridoma cells expressing the paired alpha and beta chains identified in the pancreas to test their reactivity to the cognate peptide reported in IEDB. We analyzed all TCRs from donors with a matched HLA-I allele and available counter-paired alpha or beta chain sequences (**Supplementary Table 5**). Fourteen of 15 TCR clonotypes reported to be specific for influenza A virus and two of two clonotypes reported to be specific for Epstein-Barr virus (EBV) were confirmed to respond to the cognate peptide, demonstrating accurate prediction of TCR specificity for influenza A virus and EBV, whereas none of the TCRs reported to recognize other antigens responded (**Figure 2C**, **Supplementary Table 6, Supplementary Figure 2**). The presence of influenza A-reactive T cells in islets was further supported by an independent study showing that T cell clones established from the islets of one T1D donor, nPOD6472, responded to the same influenza A peptide (**Supplementary Figure 3**). Altogether, searching a public TCR database successfully identified influenza A virus- and EBV-reactive TCR clonotypes in the pancreas of both T1D and non-diabetic donors, while identification of other antigen-reactive TCRs remained limited.

### Virus-reactive T cells are present in the pancreas of donors with and without T1D

Given the identification of TCR clonotypes shared with those reactive to influenza A- and EBV-derived peptides curated in the public TCR database, we further expanded our search to additional viral organisms, including Coxsackievirus B, which has been suggested to be associated with T1D development (*26–28*). We analyzed a total of 542 TCR clonotypes expressed by CD8 T cells in the pancreas of 25 organ donors with and without T1D (**Supplementary Table 1**) and tested their responses to 30 peptides derived from viruses, including influenza A virus, EBV, Coxsackievirus B, and others, in the presence of APCs expressing autologous HLA-I molecules. These peptides were selected based on previous evidence of T cell recognition or presentation by HLA-I molecules (**Table 2**).

**Table 2.**
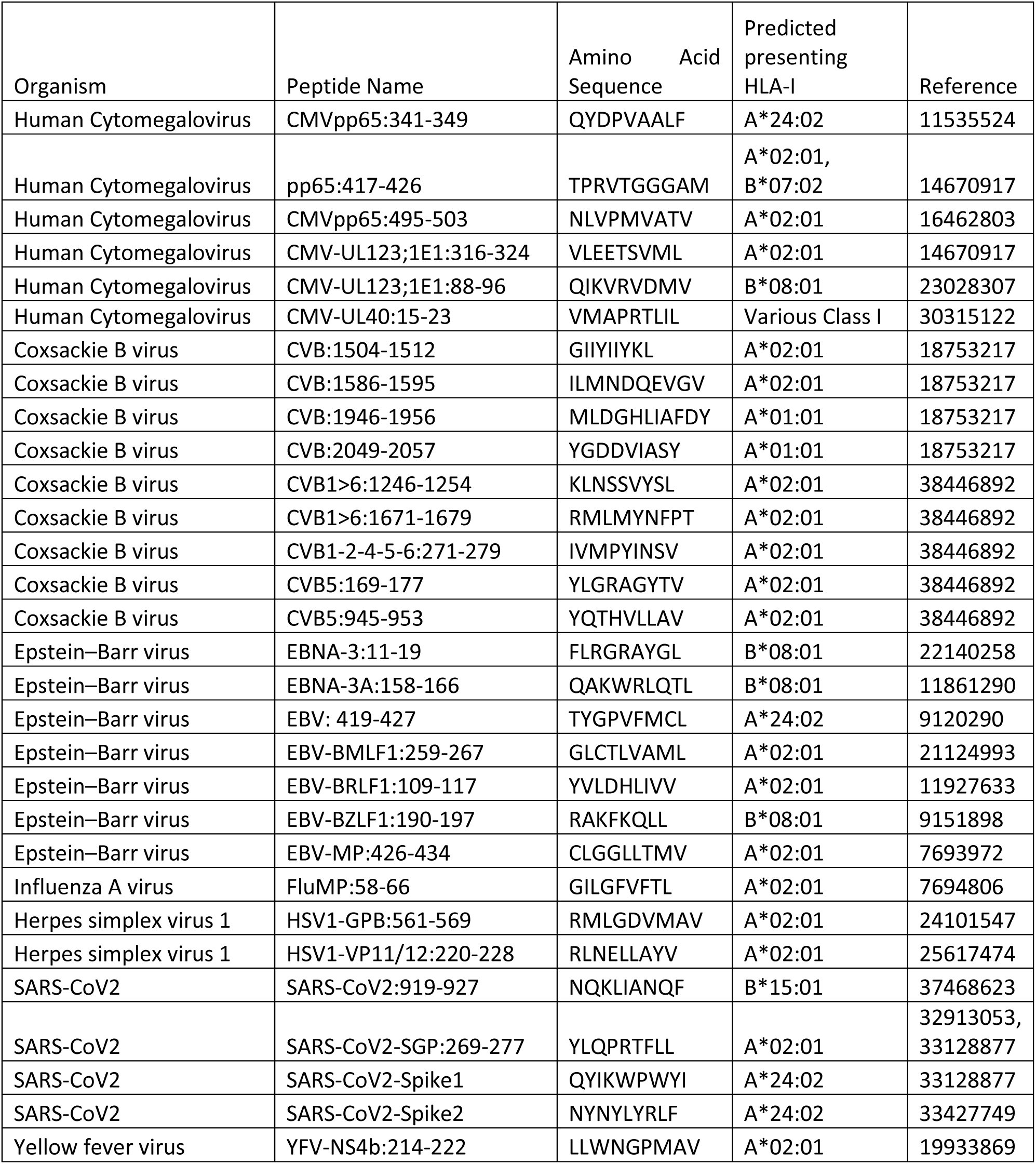
Viral peptides examined for T cell responses.

Among 542 TCRs analyzed, eleven were reactive to a peptide from influenza A virus, including eight already identified in the public TCR database search described above, and two exhibited cross-reactivity to multiple epitopes, including one responding to PPI and EBV (231.A4) and another (231.C5) to EBV and cytomegalovirus peptides (**Supplementary Figure 4, Supplementary Table 6 and 7**). On the other hand, no TCRs responded to peptides derived from Coxsackievirus B or other tested viruses (**Supplementary Table 7**). Overall, when including all TCR clonotypes identified through the public TCR database search, CD8 T cells reactive to influenza A virus- and EBV-derived peptides were detected in approximately one third and 10% of donors, respectively (**Figure 2D**). The frequencies of these viral peptide-reactive T cells were typically a few percent of CD8 T cells analyzed for the specificity (**Figure 2E, Supplementary Table 7**). Importantly, there was no significant difference between donors with and without T1D in the frequency of CD8 T cells reactive to the studied viral peptides in the pancreas (**Figure 2D and 2E**). In sum, screening hundreds of TCRs expressed by pancreatic CD8 T cells identified TCRs reactive to influenza A virus- and EBV-derived peptides, but we did not find any CD8 T cells reactive to the tested peptides derived from Coxsackievirus B in the pancreas of any of the 25 donors studied.

### Identifying TCR clonotypes reactive to stem cell-derived beta cells

Because virus-reactive T cells are present in the pancreas, we next asked whether tissue specificity is required for CD8 T cells to migrate to the islets and how frequently tissue-specific T cells are present in the pancreas with or without beta cell autoimmunity. To assess reactivity to beta cells, we utilized stem cell-derived beta-like cells (sBCs) differentiated from human pluripotent stem cells (hPSCs) of a donor carrying HLA-A2 and HLA-A24 (*29*). To induce expression of HLA-I molecules, we treated sBCs with interferon-gamma and confirmed efficient HLA-I expression by flow cytometry (**Figure 3A**). Compared with K562 human myeloid cells that express only HLA-A2, sBCs expressed slightly lower levels of HLA-A2, while overall HLA-I expression detected by anti-HLA-ABC staining was higher, presumably reflecting the presence of additional HLA-I molecules other than HLA-A2.

**Figure 3.**
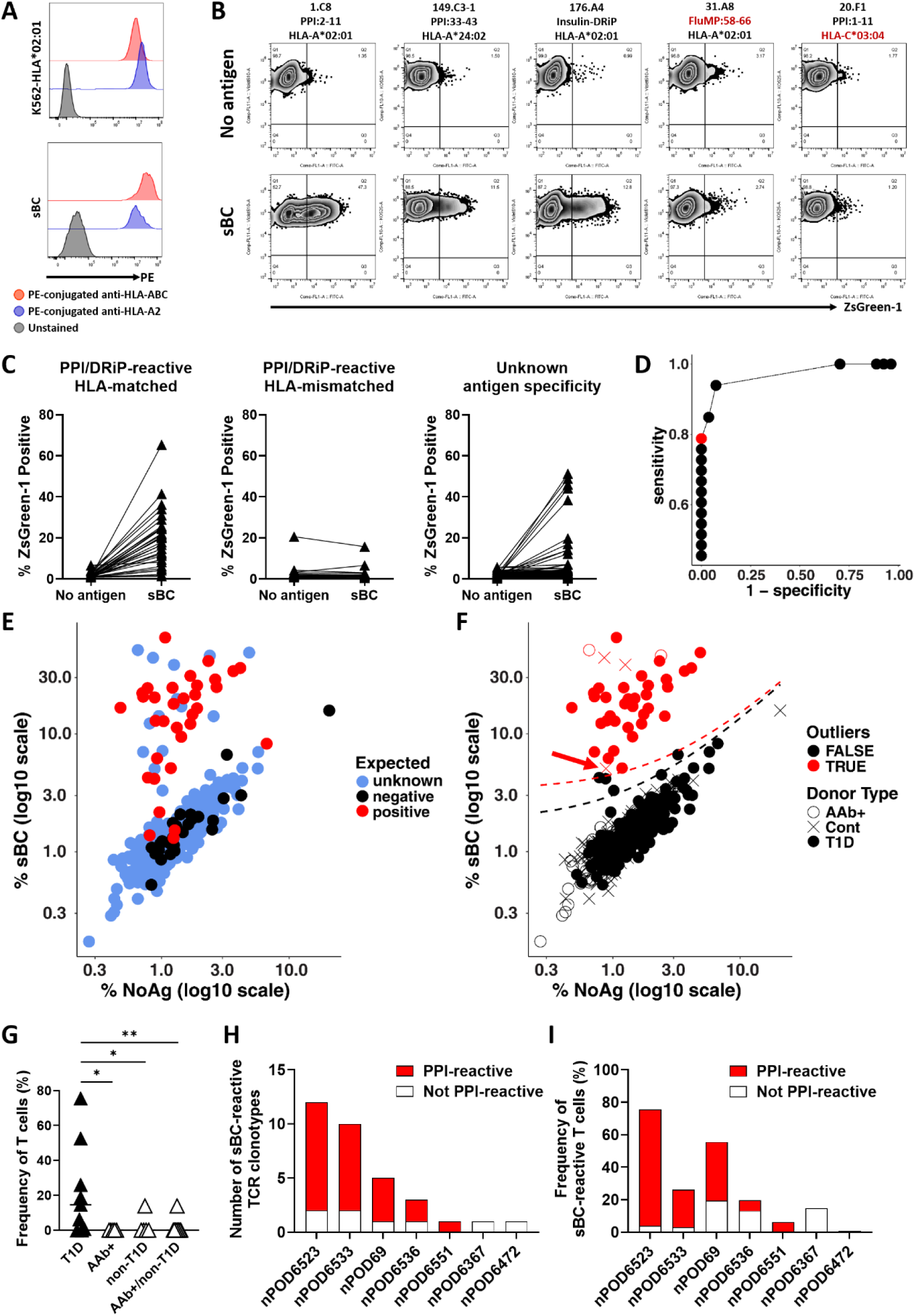
Identification of CD8 T cells reactive to stem cell–derived beta-like cells (sBCs). (A) Stem cell–derived beta-like cells (sBCs) treated with interferon-gamma and K562 cells transduced to express HLA-A*02:01 were stained with antibodies against HLA-ABC (red) or HLA-A2 (blue). Representative histograms from three independent experiments are shown. (B) TCR transductants with defined antigen reactivities (indicated above each panel) were cultured in the absence (upper panels) or presence (lower panels) of sBCs. Activation was assessed by ZsGreen1 expression driven by an NFAT reporter. Representative flow cytometry plots are shown. (C) TCR transductants were were cultured with (sBC) or without (No antigen) sBCs and were analyzed for activation determined by ZsGreen1-NFAT reporter. Percentages of ZsGreen1-positive cells with and without sBCs for each TCR clonotypes are plot by those reactive to islet peptides restricted with HLA-A2 or HLA-A24, which are expressed by sBCs (HLA-matched), those reactive to islet peptides restricted with HLA-I molecules other than HLA-A2 and HLA-A24 (HLA-mismatched), and those with unknown specificity (Target unknown). (C) TCR transductants were cultured with (sBC) or without antigen (No antigen), and activation was measured by ZsGreen1 expression. Percentages of ZsGreen1-positive cells are shown for individual TCR clonotypes grouped as those reactive to islet peptides presented by HLA-A2 or HLA-A24 (HLA-matched to the sBC donor), those reactive to islet peptides restricted by other HLA-I molecules (HLA-mismatched), or those with unknown specificity. (D) Receiver operating characteristic (ROC)-like analysis was used to define thresholds for sBC reactivity based on residuals from regression of responses in the presence versus absence of sBCs. Sensitivity is plotted against 1 − specificity across tested thresholds, and the selected cutoff is indicated by a red symbol. (E) Responses of individual TCR clonotypes are plotted as mean ZsGreen1-positive frequencies in the presence (sBC) versus absence (No antigen) of sBCs (log10 scale). TCRs are colored based on expected reactivity (positive: HLA-matched to the sBC donor; negative: HLA-mismatched; unknown: target unknown). (F) Identification of sBC-reactive TCR clonotypes based on regression analysis. TCRs above the defined threshold are classified as reactive (red), and those below as non-reactive (black). Dashed lines indicate the fitted regression (black) and the threshold used to define reactivity (red). Donor origin is indicated by symbol shape (black circles: T1D; white circles: autoantibody-positive; crosses: non-diabetic controls). (G) Frequencies of CD8 T cells expressing sBC-reactive TCRs are shown for individual donors. Each point represents the proportion of sBC-reactive T cells among all T cells expressing tested TCRs in that donor. Frequencies across donor groups (T1D, autoantibody-positive, and non-diabetic controls) were compared using the Mann–Whitney test. *p < 0.05, **p < 0.005. (H) Numbers of sBC-reactive TCR clonotypes identified for each donor. Among sBC-reactive clonotypes, red and white indicate clonotypes that are reactive or non-reactive to PPI, respectively. (I) Frequencies of CD8 T cells expressing sBC-reactive TCRs stratified by PPI reactivity (red: PPI-reactive; white: non–PPI-reactive) are shown for individual donors. Each bar represents the proportion of sBC-reactive T cells among all T cells expressing tested TCRs in that donor.

To evaluate the specificity of TCR responses to sBCs, we analyzed reactivity using representative TCRs whose antigen specificity and HLA restriction had already been identified. TCRs reactive to PPI or insulin-DRiP peptides presented by either HLA-A2 or HLA-A24 responded to sBCs. In contrast, a TCR reactive to an influenza viral peptide presented by HLA-A2 and a TCR reactive to a PPI peptide presented by HLA-C*03:04, an allele not carried by the sBC donor, did not (**Figure 3B**). These results indicate that TCR responses to sBCs require both islet antigen and matched HLA-I molecules.

### Beta cell-reactive TCR clonotypes are found in the pancreas of T1D but are rare in non-diabetic organ donors

Using sBCs as targets, we analyzed the responses of 426 TCRs expressed by CD8 T cells in the pancreas of donors carrying HLA-A2 or HLA-A24, including those identified as virus-reactive (**Supplementary Table 1**). This cohort comprised nine donors with T1D, four donors positive for anti-GAD antibodies, and six non-diabetic control donors. In addition to HLA-A2 and HLA-A24, a few GAD antibody-positive and non-diabetic donors shared HLA-B or HLA-C alleles with the sBC donor (**Supplementary Table 8)**. As expected, many PPI- and insulin-DRiP-reactive TCRs restricted by HLA-A2 or HLA-A24 responded to sBCs, whereas those restricted by other HLA-I molecules did not (**Figure 3C**). Of note, all the PPI- and insulin-DRiP-reactive TCRs restricted by HLA alleles other than HLA-A2 and HLA-A24 are restricted by HLA alleles not carried by the sBC donor; thus, their HLA restrictions are mismatched to those expressed by sBCs. Notably, several TCRs of unknown antigen specificity also responded to sBCs.

To evaluate the performance of the sBC assay, we modeled responses using a regression framework with islet peptide-reactive TCRs restricted by HLA-A2 or HLA-A24 as expected positives and those restricted by other HLA-I molecules as expected negatives (**Figure 3D and 3E**). The assay efficiently identified TCR clonotypes reactive to sBCs, yielding an AUC value of 0.97 (**Figure 3D**), confirming that sBCs express the relevant epitopes presented by HLA-I molecules recognized by islet peptide-reactive TCRs. Using a cutoff that ensured 100% specificity while achieving the highest sensitivity, which detects 79% of islet peptide-reactive TCR clonotypes restricted by HLA-A2 or HLA-A24, we classified 39 of the 426 TCR clonotypes, including 13 with unknown antigen specificity, as sBC-reactive (**Figure 3E and 3F**). To exclude allogeneic responses, we tested whether these 13 antigen-unknown TCRs were reactive to K562 cells expressing each HLA-I molecule of the sBC donor. Four of five TCRs derived from non-diabetic donors, but only one of eight from T1D donors, were classified as reactive to the sBC donor’s HLA-I molecules and considered alloreactive (**Supplementary Figure 5**). The remaining TCRs are likely to be truly reactive to epitopes uniquely expressed by sBC.

After excluding alloreactive clonotypes, 33 out of 245 TCRs from T1D donors were reactive to sBCs expressing HLA-A2 and HLA-A24, whereas only one of 181 TCRs from GAD antibody-positive or non-diabetic donors showed minimal reactivity (p<0.0001, Chi-square test). Of note, this single TCR from a non-diabetic donor exhibited barely detectable reactivity, as indicated by the arrow in **Figure 3F**. When analyzing frequencies of CD8 T cells expressing sBC-reactive TCRs, T1D donors contained significantly higher proportions of sBC-reactive T cells in the pancreas than non-T1D or GAD antibody-positive donors (**Figure 3G**, **Table 1, Supplementary Table 8**). Thus, the presence of CD8 T cells expressing TCRs highly reactive to beta cells is a defining feature in the pancreas of T1D donors. Importantly, none of the virus-reactive TCRs (n=20) responded to sBCs, regardless of the disease status, indicating that their presence is not attributed to cross-reactivity with antigens expressed by HLA-I molecules on the cell surface of sBCs. Together, these findings indicate that the entry of CD8 T cells highly reactive to beta cells into the pancreas is selectively permitted in T1D donors but tightly forbidden in donors without T1D, while, at the same time, tissue specificity is not necessarily required for their entry.

### Native preproinsulin 15-24 serves as a biological epitope recognized by beta cell-reactive TCRs

Among sBC-reactive T cells in the pancreas of T1D donors, 25 of 33 (76%) TCR clonotypes and 157 of 195 (80%) T cells expressing sBC-reactive TCRs were PPI-reactive, although this proportion varied across individual donors (**Figure 3H and 3I, Supplementary Table 8**). To dissect the relationship between TCR responses to beta cells and PPI, we focused on TCR clonotypes reactive to PPI:15-24 within the beta cell-reactive repertoire, as this epitope is dominantly presented by HLA-I molecules on pancreatic beta cells, as demonstrated by immunopeptidome analyses (*30–33*). If PPI:15-24 functions as a biological epitope that activates CD8 T cells in the pancreas, TCR responsiveness to beta cells should correlate with responsiveness to this peptide. To test this hypothesis, we compared the magnitude of responses to PPI:15-24 and to beta cells using PPI:15-24-reactive TCR transductants. Seven TCR clonotypes responded to PPI:15-24 presented by HLA-A2-expressing K562 cells at varying levels (**Figure 4A**). To minimize potential allogeneic responses to minor donor-specific antigens, we assessed reactivity to primary islets isolated from three independent donors. One TCR, 1.F1, was identified as an outlier based on Cook’s distance (2.51 > 0.571). After excluding this outlier, a significant positive correlation was observed between responses to PPI:15-24 and human islets (**Figure 4B**, Pearson’s r=0.88, p=0.021). The disproportionately strong response of 1.F1 to human islets may reflect cross-reactivity to other epitopes, including post-translationally modified (PTM) forms of PPI:15-24. To explore this possibility, we examined responses to three PTM versions of PPI:15-24: two detected by immunopeptidome analyses (N-terminal acetylation of alanine at position 24, and substitution of tryptophan with hydroxytryptophan at position 17) (*30, 31*) and one predicted modification reported in tumors (substitution of tryptophan with phenylalanine at position 17) (*34*). All seven TCR clonotypes, including 1.F1, responded more strongly to the native PPI:15-24 than to any of the PTM variants (**Figure 4C**). Together, these findings indicate that all TCRs except 1.F1 recognize human islets through the native PPI:15-24 epitope, supporting the conclusion that the unmodified PPI:15-24 peptide serves as a biological epitope for the majority of PPI:15-24-reactive TCR clonotypes.

**Figure 4.**
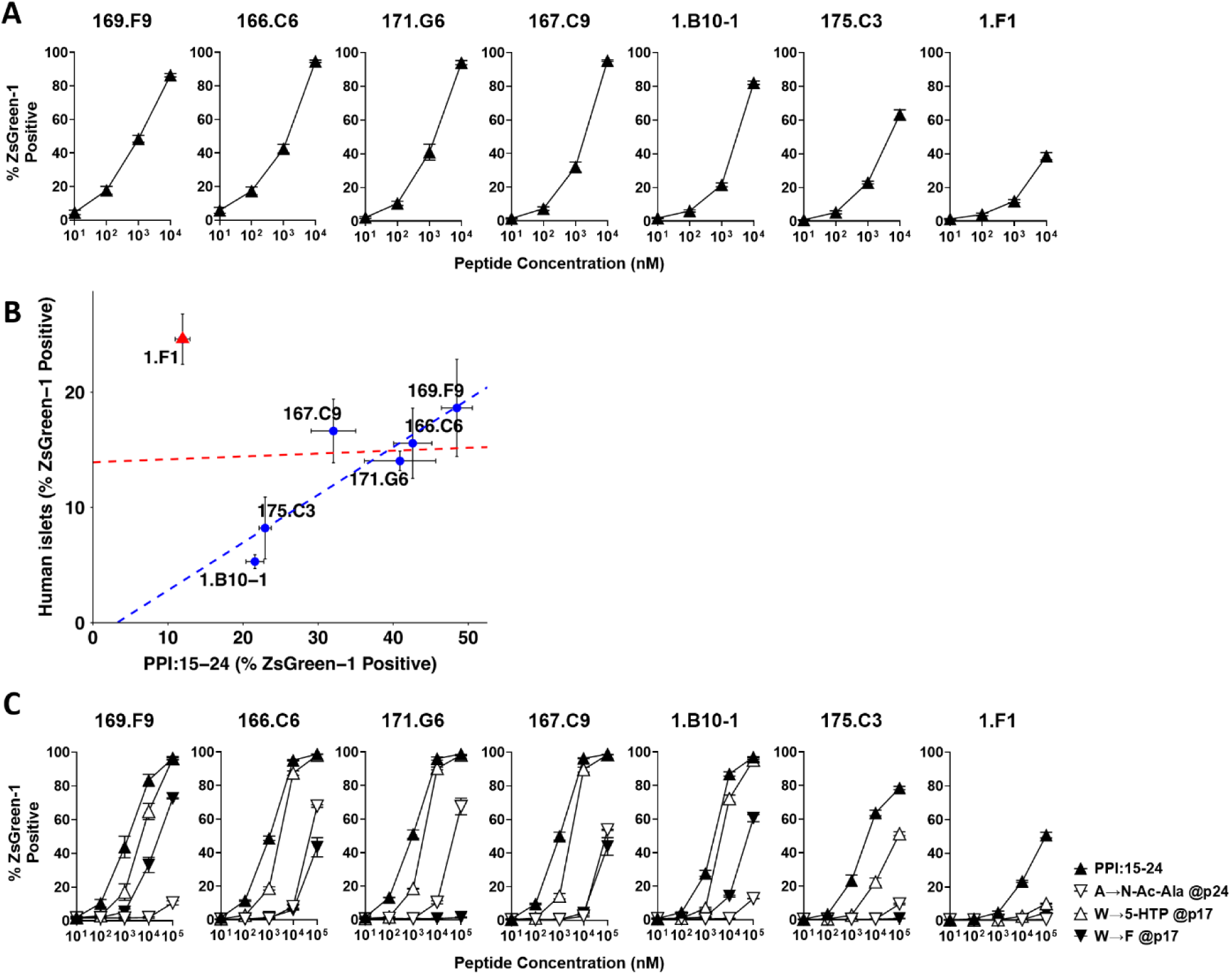
Reactivity to PPI peptide and sBCs. (A) PI:15–24–reactive TCR transductants were tested for responses to increasing concentrations of the PPI:15–24 peptide in the presence of K562 cells expressing HLA-A*02:01. Activation was assessed by ZsGreen1 expression driven by an NFAT reporter, and mean values ± SEM from five independent experiments are shown. (B) Responses to sBCs (mean ± SEM from two independent experiments) are plotted against responses to PPI:15–24 peptide stimulation (1 µM; mean ± SEM from five independent experiments). Red and blue dashed lines indicate correlations across all TCR clonotypes and with exclusion of TCR 1.F1, respectively. (C) PPI:15–24–reactive TCR transductants were tested for responses to increasing concentrations of modified PPI:15–24 peptides in the presence of K562 cells expressing HLA-A*02:01. Activation was assessed by ZsGreen1 expression driven by an NFAT reporter, and mean values ± SEM from two independent experiments are shown.

## Discussion

Analysis of TCRs expressed by pancreatic resident CD8 T cells revealed that T cells highly reactive to beta cells are present in the pancreas of individuals with T1D but are largely absent in non-diabetic donors. In contrast, virus-reactive CD8 T cells were detected in the pancreas of both T1D and non-diabetic donors, indicating that tissue residency alone does not confer beta cell specificity. These findings demonstrate that the presence of beta cell-reactive CD8 T cells, rather than general T cell infiltration, is a defining feature of the T1D pancreas. The selective presence of beta cell-reactive T cells in the T1D pancreas supports their involvement in disease development and aligns with the targeted destruction of beta cells, indicating that tolerance among T cells recognizing beta cell-derived epitopes is broken in T1D. Notably, reactivity to native peptides derived from major islet proteins other than PPI was rarely detected, suggesting that tolerance to these antigens remains largely intact even at the stage of clinical T1D, while T cells recognizing additional beta cell-derived epitopes are present. As beta cell reactivity among T1D-associated T cells represents a robust functional hallmark of disease, this property, independent of precise epitope definition, may provide a practical framework for identifying pathogenic T cells and for developing biomarkers and targeted immunotherapies.

Defining the proportion of tissue-reactive T cells is critical for understanding their contribution to pathogenicity and for identifying which T cell populations must be targeted to halt beta cell autoimmunity. In this context, assays measuring responses to sBCs provide a functional and reproducible approach to detect beta cell reactivity among pancreatic CD8 T cells. Although the current platform does not capture all possible HLA-I-restricted responses, as the sBCs used express only a single HLA-I (HLA-A2 or HLA-A24) shared with individual T1D donors, it enables an initial estimate of the frequency of beta cell-reactive T cells. Despite this limitation, five of seven donors exhibited frequencies approaching or exceeding ∼1/6 of CD8 T cells reactive to sBCs (**Figure 3I**), suggesting that a substantial fraction of pancreatic CD8 T cells in T1D may be beta cell-reactive. This estimate is consistent with prior observations that CD8 T cells binding to PPI:15-24 presented by HLA-A2 can account for approximately 10% of pancreatic CD8 T cells in T1D donors (*16*). Future studies incorporating sBCs expressing a broader range of HLA-I molecules will enable more precise quantification of tissue-reactive T cells. Notably, recent evidence that HLA-B expression is enhanced by interferon-alpha suggests that CD8 T cells restricted by HLA-B may contribute significantly to disease pathogenesis (*31*). Accordingly, engineering sBC platforms to express individual HLA-I molecules, including HLA-B and HLA-C, matched to donor haplotypes will be important to fully define the landscape of beta cell-reactive T cells in the pancreas.

While beta cell reactivity is a defining feature of CD8 T cells in the pancreas of T1D donors, virus-reactive T cells are also present, consistent with prior studies demonstrating their detection in pancreatic tissues by tetramer-based histological analysis (*16*). The absence of Coxsackievirus B-reactive CD8 T cells in islets is consistent with a previous study, which detected such T cells in the spleen and pancreatic lymph nodes, but not in the pancreas, using in situ HLA-I multimer staining (*35*). Because virus-reactive T cells are observed in both T1D and non-diabetic donors, their presence in the pancreas likely reflects a physiological feature independent of disease status. However, their presence under physiological conditions does not exclude a potential role in beta cell autoimmunity, either by promoting or modulating immune responses (*36*). In the current study, none of the virus-reactive TCRs tested responded to sBCs in our assay system, which robustly detects beta cell reactivity, with nearly 80% of PPI-reactive TCRs showing HLA-restricted responses (**Figure 3D**), providing no indication of detectable cross-reactivity between the viral epitopes examined and beta cell antigens under the conditions tested. Notably, studies in mouse models have suggested that non-islet-reactive CD8 T cells infiltrate pancreatic islets and can influence disease progression by limiting access of islet-reactive T cells to target tissues, thereby suppressing their activation and expansion (*37, 38*). Whether virus-reactive CD8 T cells in humans exert similar modulatory effects remains an open question. Addressing this issue will be critical for understanding the complex role of viral infections in T1D pathogenesis.

Focusing on the antigen specificity of beta cell-reactive T cells, a substantial fraction of sBC-reactive CD8 T cells recognized PPI (**Figure 3I**), yet the diversity of epitopes was strikingly limited. Although PPI comprises 110 amino acids, analysis of eight T1D donors carrying HLA-A*02:01, including those reported previously (*17*), revealed that all 23 TCRs reactive to PPI peptides presented by HLA-A*02:01 recognized one of two regions: PPI:1-11 through 3-13 or PPI:15-23 through 15-25. An additional epitope derived from insulin-DRiP (1-9 through 1-11) has also been identified in the context of HLA-A*02:01, with no evidence for broader epitope distribution across the molecule. Similarly, among TCRs restricted by HLA-C*03:04, 17 clonotypes identified from three of five T1D donors recognized only three epitopes (PPI:1–11, PPI:23–30, and PPI:69–77), which, together with an additional region (PPI:100-110) identified in another comprehensive analysis recently reported (*25*), define a limited set of targeted regions within PPI, further underscoring the restricted epitope landscape. These data indicate that CD8 T cell autoreactivity in T1D is directed toward a highly restricted set of epitopes within PPI, supporting a model in which pathogenic responses are driven by focused antigen recognition rather than broad targeting of the islet proteome. This pattern is consistent with CD8 T cell responses to viral infection, which are typically focused on a limited number of immunodominant epitopes (*39, 40*). Thus, our data demonstrate that CD8 T cell responses in T1D are tightly constrained and focused across individuals sharing the same HLA allele, paralleling immunodominance patterns described in viral infection. Although a subset of sBC-reactive T cells remains with unknown antigen specificity, it is plausible that their target epitopes may be similarly constrained. In this context, immunopeptidomics analyses have provided valuable insight into peptides presented by HLA-I molecules in islets (*30–33*), enabling the identification of candidate antigens for CD8 T cells in T1D. A major challenge, however, lies in narrowing thousands of eluted peptides to a tractable set of functionally relevant epitopes. Emerging high-throughput screening platforms offer powerful approaches to address this problem (*41–44*), and their integration with immunopeptidomics and mass spectrometry is likely to accelerate the identification of antigens recognized by T cells with currently unknown specificity.

The limited detection of TCRs reactive to native peptides from islet proteins other than PPI was unexpected. Previous studies have identified CD8 T cells specific for IGRP and ZnT8 peptides using fluorescence-conjugated multimers in histological analyses of pancreatic tissues (*13, 15*). Several explanations may account for this discrepancy. First, these T cells may recognize islet-derived peptides with relatively low affinity and/or exhibit promiscuous antigen recognition, resulting in weak or indistinct functional responses at the clonal level. Indeed, individual T cells can recognize a large number of distinct epitopes, including both self- and pathogen-derived peptides (*45, 46*), and cross-reactive CD4 T cell clones recognizing multiple islet antigens have been described in T1D (*47*). Consistent with this, two of the three TCRs identified in our study as reactive to major islet protein-derived peptides exhibited promiscuous recognition of multiple epitopes. These observations raise the possibility that some pancreatic T cells bind multiple peptide-HLA complexes, including those derived from major islet proteins, and can therefore be detected by multimer staining, but do not generate sufficiently strong signaling to elicit measurable NFAT-dependent activation in functional assays. Second, TCRs analyzed in the current study were selected from clonotypes expressed by multiple cells. T cells reactive to native islet peptides may not undergo substantial clonal expansion and therefore may be underrepresented in our dataset. If so, this would further suggest that CD8 T cells reactive to these native islet peptides are relatively infrequent in the pancreas. Future studies analyzing antigen specificity across the full TCR repertoire at the single-clonotype level will help clarify this possibility. Third, CD8 T cells reactive to native islet peptides may not be stably retained in the pancreas, potentially due to low-affinity or promiscuous interactions with antigen and may instead be more readily detected in peripheral circulation or pancreatic lymph nodes. Consistent with this, several studies have reported the presence of islet peptide-reactive CD8 T cells in peripheral blood, including reduced frequencies of GAD- and ZnT8-reactive T cells following anti-CD3 monoclonal antibody therapy (*15, 48–50*). Thus, although these cells appear to be infrequent in the pancreas, they may still have utility as peripheral biomarkers.

Beyond these considerations, analysis of TCRs for reactivity to sBCs identified a subset of beta cell-reactive TCRs with currently unknown antigen specificity. Because these TCRs did not respond to native peptides derived from major islet proteins (GAD65, ZnT8, IGRP, ChgA, and IAPP), their targets may include peptides derived from other islet proteins not examined in this study, such as secretogranin 5 (SCG5) and insulinoma-associated protein 2 (IA-2), or neoepitopes, including PTM or RNA splice variant-derived peptides. Regardless of their precise identity, it is critical to distinguish “biological epitopes,” which elicit functional T cell responses in vivo, from “mimotopes,” which can bind TCRs but are not relevant to beta cell destruction, as these repertoires only partially overlap (*4, 51*). Increasing evidence indicates that T cells reactive to PTM self-antigens contribute to T1D pathogenesis (*7, 32, 52–58*). Notably, native and PTM antigens often share core sequence motifs, enabling cross-reactivity. For example, CD4 T cells isolated from the pancreas of T1D donors have been shown to respond to both C-peptide and hybrid insulin peptides (HIPs), a major class of PTM antigens in T1D (*59*). This complexity highlights a key challenge in defining antigen specificity: responses to peptides do not necessarily predict reactivity to beta cells. Indeed, responses to beta cells and to peptides were generally correlated for PPI:15–24-reactive TCRs, with notable exceptions (**Figure 4B**), indicating that peptide reactivity alone may not fully predict beta cell recognition. These findings underscore the importance of directly assessing T cell reactivity to beta cells to identify antigens that truly drive functional responses. In this context, because neoepitopes may be generated preferentially under disease-associated conditions (*60–65*), evaluating T cell responses to beta cells exposed to relevant stressors will be important. While the current study utilized interferon-gamma to induce HLA-I expression, future studies incorporating additional conditions, such as interferon-alpha or ER stress (e.g., tunicamycin), will help define the spectrum of epitopes that activate pathogenic T cells and contribute to the break in tolerance in T1D.

In conclusion, our analysis of pancreatic CD8 TCR repertoires defines beta cell reactivity as a distinguishing functional property of T cells associated with T1D, rather than a general feature of tissue-resident T cells. These cells are present at substantial frequencies in the T1D pancreas and exhibit a highly restricted antigenic landscape. These findings support a model in which T1D is driven by a focused CD8 T cell response to beta cells, providing a framework for understanding disease mechanisms and a rationale for leveraging beta cell reactivity to identify T1D-associated T cells for biomarker development and targeted therapies.

## Methods and Materials

### Study approvals

Pancreas slice tissues and pancreatic islets were obtained from the Network for Pancreatic Organ Donors with Diabetes program (nPOD; RRID:SCR_014641) (*66*), the Integrated Islet Distribution Program (IIDP; RRID:SCR_014387), and the Alberta Diabetes Institute IsletCore (ADI). For pancreas slice samples obtained through nPOD, pancreatic islets were further isolated by enzymatic digestion with Liberase DL (Roche). Measurement of type 1 diabetes–associated autoantibodies and HLA typing for nPOD donors were performed according to nPOD Standard Operating Procedures (https://www.jdrfnpod.org/for-investigators/standard-operating-procedures/). Use of these postmortem tissues was reviewed by the Colorado Multiple Institutional Review Board and approved as Not Human Subjects Research.

### Donor information and TCR sequencing

Information on pancreas and islet donors, including those reported previously (*17*), is provided in **Supplementary Table 1**. T cells were isolated from islet or pancreas slice samples, and TCR alpha and beta chain genes were amplified from single cells using a multi-step PCR approach as described previously (2), using modified primers. Illumina adapter and index sequences were introduced during later amplification steps, with indexing performed at the final stage for MiSeq libraries and across multiple amplification steps for NovaSeq libraries (**Supplementary Table 9**). TCR amplicons were sequenced on MiSeq or NovaSeq platforms (Illumina), as indicated in **Supplementary Table 1**. Sequence reads were demultiplexed using a custom Python script [https://github.com/CUAnschutzBDC/pancreatic_cd8_t_cell_repertoire] and analyzed using IMGT/HighV-QUEST (https://www.imgt.org/HighV-QUEST/). IMGT output files were further processed using a custom Python script [https://github.com/CUAnschutzBDC/pancreatic_cd8_t_cell_repertoire] to assign variable and joining genes and define TCR junction sequences for individual cells. The annotations were subsequently reviewed and manually curated to remove low-frequency sequences that represented a minor fraction of reads within individual cells and were considered likely artifacts based on their abundance and comparison with negative-control wells, in which no cells were sorted.

### Generation of TCR and HLA transductants and EBV-transformed autologous B cells

TCR- and HLA-expressing transductants were generated using murine T cell–derived 5KC T-hybridoma cells engineered with a ZsGreen1 NFAT reporter and human myeloma–derived K562 cells, respectively, as previously described (*17, 67, 68*). HLA allele sequence information was obtained from the IPD-IMGT/HLA database (https://www.ebi.ac.uk/ipd/imgt/hla/). Autologous EBV-transformed B cell lines were established from splenocytes of individual islet donors using standard methods (*69*).

### Screening for reactivity to islet and viral peptides

A multiplex T cell stimulation assay was used to assess reactivity to truncated peptide pools and individual viral peptides, as previously described (*17, 67*). Briefly, up to eight T-hybridoma transductants expressing TCRs derived from each individual donor were mixed and cultured (20,000 cells/well per clonotype) with or without antigen in the presence of antigen-presenting cells. Autologous EBV-transformed B cells (100,000 cells/well) or a mixture of K562 cells expressing the HLA-A, HLA-B, and HLA-C alleles of the corresponding donor (105,000 cells/well) were used as antigen-presenting cells, and the type of antigen-presenting cells used for each donor is indicated in **Supplementary Table 1**. Peptide pools consisting of 8–11-mer crude peptides sharing a common C-terminal residue at each position within PPI, insulin DRiP, GAD65, IAPP, ChgA, ZnT8, and IGRP were custom-ordered from Mimotopes and added at a total peptide concentration of approximately 200 µg/ml. The amino acid sequences included in each pool are provided in **Supplementary Table 10**. Individual viral peptides (**Table 2**), synthesized by Genemed Synthesis or GenScript, were used at 100 µM for screening. Cultures with 5 µg/ml of anti-mouse CD3ε antibody (clone 125-2C11, BD) were included in the assay as a positive control. After overnight stimulation, ZsGreen1 expression in each TCR transductant was quantified by flow cytometry using a CytoFLEX (Beckman Coulter) or Quanteon (Agilent) instrument and analyzed using FlowJo software (BD Biosciences). To normalize background activation (i.e., the percentage of ZsGreen1-positive cells in cultures without antigen), responses to each peptide or peptide pool were calculated for each TCR clonotype as the percentage of ZsGreen1-positive cells in antigen-stimulated cultures minus the average percentage of ZsGreen1-positive cells across all antigen conditions for that TCR clonotype within the same assay. Responses with normalized ZsGreen1-positive values >30% were classified as positive. This threshold was empirically determined based on the reproducibility of positive responses across replicate stimulations with synthetic peptides of ≥85% purity.

### Identification of HLA restriction and titration analysis

TCR transductants that responded to islet peptide pools or viral peptides were cultured (20,000 cells/well) with or without the cognate peptide pool at a total peptide concentration of 200 µg/ml for PPI and insulin-DRiP screening, individual peptides (100 µM) contained in the cognate peptide pool for other islet protein screening, the cognate viral peptide (100 µM), or anti-mouse CD3ε antibody (clone 125-2C11, BD, 5 µg/ml) in the presence of K562 cells expressing each HLA-I molecule carried by the TCR donor (50,000 cells/well) to determine the HLA molecule presenting the peptide to the TCR, and ZsGreen1 expression was evaluated by flow cytometry. For TCRs that responded to multiple islet peptide pools, the peptide pool that stimulated the TCR transductants most strongly was used to determine HLA restriction. For titration analysis, peptides contained in islet peptide pools that stimulated TCR transductants were synthesized by Genemed Synthesis or GenScript. Once the restricting HLA was determined, individual TCR transductants (20,000 cells/well) were cultured with peptides at concentrations ranging from 10 pM to 100 µM in the presence of HLA transductants expressing the cognate HLA-I molecule (50,000 cells/well) overnight, followed by evaluation of ZsGreen1 expression by flow cytometry. EC_50_ values were calculated using GraphPad Prism using a nonlinear regression model (log[agonist] vs. response, three parameters).

### Public TCR database search

We utilized the Immune Epitope Database & Tools (IEDB, https://www.iedb.org/) to search for TCR sequence information reactive to known epitopes. TCR sequence and epitope information were downloaded from IEDB on April 16, 2025, using the following criteria: (1) epitopes included linear peptides; (2) TCRs and epitopes were determined by T cell and MHC ligand assays; (3) TCRs were restricted by HLA-I molecules; and (4) TCRs were derived from human hosts. TCR alpha or beta clonotypes sharing the same V-gene, J-gene, and exact junction amino acid sequence as publicly reported TCRs were identified among islet- and pancreas-derived TCRs using a custom Python script [https://github.com/CUAnschutzBDC/pancreatic_cd8_t_cell_repertoire]. To maximize the accuracy of public TCR identification, only TCRs that were reported in two or more independent publications were classified as public TCRs.

### T cell clone peptide-stimulation assay

Single islets were hand-picked from enzyme-dispersed live slices of pancreas from nPOD donor 6472 as described previously (*32*). Islet-derived T cell lines were tested for influenza A peptide GILGFVFTL reactivity and reactive lines were cloned by standard 0.3 cell/well limiting dilution with irradiated allogeneic PBMC and 4.5 ug/mL PHA-P (Sigma) with additions of IL-2 and IL-15 (*32*). Expanded clones were tested for reactivity to influenza A peptide GILGFVFTL. Reactive T cell clones (25,000 cells/well) were cultured with titrated concentrations of influenza A peptide GILGFVFTL or the control PRAME_425-433_ peptide SLLQHLIGL, ranging from 8 nM to 10 µM, in the presence of the irradiated human B-cell lymphoblastoid C1R line expressing HLA-A2 (10,000 cells/well) in round-bottom 96-well plates. After 48–72 h of culture, supernatants were collected, and interferon-gamma secretion was measured by standard ELISA according to the manufacturer’s instructions (BD Biosciences).

### Generation of stem cell-derived beta like cells

Stem cell-derived beta like cells (sBCs) were generated from a human pluripotent stem cells (hPSC) Mel1^INS-GFP^ line [National Institutes of Health (NIH) registry #0139] (*29*) using a suspension-based, magnetic stirring system (REPROCELL) as previously described (*70, 71*). Briefly, Mel1 line was maintained in 2D adherent culture as colonies on human embryonic stem cell qualified Cultrex (Biotechne) in mTeSR^+^ media (STEMCELL Technologies). To initiate suspension culture, confluent hPSC cultures were dissociated into single-cell suspensions by incubation with TrypLE (Gibco) for 6-8 min and seeded at 0.5 × 10^6^ cells/ml in mTeSR^+^ media supplemented with 10 μM ROCK inhibitor in bioreactors. After confirming three-dimensional cluster formation in 48 hours, definitive endoderm differentiation was induced using d1 media [RPMI containing 0.2% fetal bovine serum (FBS), 1:5000 insulin-transferrin-selenium (ITS) (Gibco), activin A (200 ng/ml; Bio-Techne), and 3 μM CHIR99021 (STEMCELL Technologies)]. To continue direct differentiation, following medias were used: RPMI containing 0.2% FBS, 1:2000 ITS, and activin A (100 ng/ml) at days 2 and 3; RPMI containing 2% FBS, 1:1000 ITS, and keratinocyte growth factor (KGF) (50 ng/ml; PeproTech) at days 4 and 5; Dulbecco’s modified Eagle’s medium (DMEM) with d-glucose (4.5 g/liter; Gibco) containing 1:50 N-21 MAX (Biotechne), 1:100 nonessential amino acid (NEAA; Gibco), 1 mM sodium pyruvate (Gibco), 1:100 GlutaMAX (Gibco), 3 nM 4-[(E)-2-(5,6,7,8-Tetrahydro-5,5,8,8-tetramethyl-2-naphthalenyl)-1-propenyl]benzoic acid (TTNPB) (Bio-Techne), 250 nM Sant-1 (Bio-Techne), 250 nM LDN193189 dihydrochloride (LDN) (STEMCELL Technologies), 300 nM phorbol 12-myristate 13-acetate (Sigma-Aldrich), and 50 μg/ml 2-phospho-l-ascorbic acid trisodium salt (vitamin C) (STEMCELL Technologies) at day 6; DMEM containing 1:50 N-21 MAX, 1:100 NEAA, 1 mM sodium pyruvate, 1:100 GlutaMAX, 3 nM TTNPB, and vitamin C (50 μg/ml) at day 7; DMEM containing 1:50 N-21 MAX, 1:100 NEAA, 1 mM sodium pyruvate, 1:100 GlutaMAX, epidermal growth factor (100 ng/ml; Bio-Techne), KGF (50 ng/ml), and vitamin C (50 μg/ml) at days 8 and 9; DMEM containing 1:50 N-21 MAX, 1:100 NEAA, 1 mM sodium pyruvate, 1:100 GlutaMAX, 10 μg/ml heparin (Sigma-Aldrich), 2 mM *N*-acetyl-l-cysteine (Cysteine) (Sigma-Aldrich), 10 μM zinc sulfate heptahydrate (Zinc) (Sigma-Aldrich), 1× β-Mercaptoethanol (Sigma-Aldrich), 10 μM Alk5i II RepSox (Bio-Techne), 2 μM 3,3′,5-triiodo-l-thyronine sodium salt (T3) (Sigma-Aldrich), 0.5 μM LDN, 1 μM gamma secretase inhibitor XX (XXi) (Bio-Techne), 10 μM Y-27632 dihydrochloride (selective ROCK inhibitor) (Bio-Techne), and 1:250 1 M NaOH to adjust pH to ∼7.4 at days 10 through 15; and CMRL (Gibco) containing 1:50 N-21 MAX, 1:100 NEAA, 1:100 GlutaMAX, 10 μg/ml heparin, 2 mM cysteine, 10 μM zinc, 1x BME, 2 μM T3, 10 μM Alk5i II RepSox, 50 μg/ml vitamin C, 1:1000 trace elements A (Corning), 1:1000 trace elements B (Corning), and 1:250 NaOH to adjust pH to ∼7.4 at days 16 through ∼30. All differentiation media also contained 1× penicillin-streptomycin (Sigma-Aldrich). After confirming GFP expression by flow cytometry, sBC clusters were dissociated into single cells and cryopreserved at 3 x 10^6^ cells/100 μL in CryoStor CS10 (StemCell Technologies) until being used for TCR transductant stimulation assays as described previously (*72*).

### Evaluation of responses to sBCs

Responses of TCR transductants to sBCs were evaluated using a multiplex T cell stimulation assay system (*67, 68*), in which up to 8 TCR transductants engineered with an NFAT-driven ZsGreen1 reporter were co-cultured with single cells dissociated from interferon-gamma–treated sBC clusters. Cryopreserved sBCs were thawed 1 week before use and cultured in 5 ml of sBC media (days 16–30 media described above) in six-well suspension plates at 37°C. Cells re-clustered within a few days, and interferon-gamma (PeproTech) was added at 200 ng/ml for the final 2 days of culture. sBC clusters were dissociated into single cells using 0.05% trypsin-EDTA (Gibco) at 37°C for 10–15 minutes, followed by two washes with TCR transductant culture media [IMDM (Gibco) supplemented with 10% FBS (Bio-Techne), 1× penicillin-streptomycin (Sigma-Aldrich), and 0.001% β-mercaptoethanol (Sigma-Aldrich)].

TCR transductants (12,000–20,000 cells/well per clonotype) were cultured with or without sBCs (30,000–40,000 cells/well; ∼2.5-fold excess relative to each TCR clonotype) in 200 μl of media in round-bottom 96-well plates overnight, followed by measurement of ZsGreen1 expression by flow cytometry. Anti-mouse CD3ε antibody (5 µg/ml; clone 125-2C11, BD) was included as a positive control. For assessment of HLA expression, a portion of dissociated sBCs and K562 cells transduced with HLA-A*02:01 (*17, 68*) were stained with anti–HLA-ABC (clone G46-2.6, BD) and anti–HLA-A2 (clone BB7.2, BioLegend) antibodies and analyzed on a Quanteon flow cytometer (Agilent).

### Regression model to define sBC-reactive TCR clonotypes

TCR reactivity was quantified by comparing responses in the presence and absence of sBCs. For each TCR, replicate %ZsGreen1-positive measurements were averaged, and a linear regression model was fit to mean sBC responses as a function of mean responses without antigen. Residuals from this model were used to identify TCRs with elevated sBC reactivity relative to background. To define an objective threshold for positivity, a range of cutoffs was evaluated by applying multiples of the residual standard deviation, and corresponding sensitivity and specificity values were calculated based on expected reactivity annotations. Receiver operating characteristic (ROC)-like curves were generated by plotting sensitivity against 1 − specificity across thresholds. The optimal threshold was selected as the most stringent cutoff (highest standard deviation multiplier) that maintained perfect specificity while maximizing sensitivity. The area under the ROC curve (AUC) was calculated using the trapezoidal rule to summarize overall classification performance. All analyses and visualizations were performed in R using custom scripts [https://github.com/CUAnschutzBDC/pancreatic_cd8_t_cell_repertoire].

### Evaluation of response to allogeneic HLA molecules

The Mel1 sBC donor carried HLA-A*02:01, A*24:02, B*07:02, B*14:01, C*02:02, and C*08:02. K562 expressing each of these HLA alleles were generated as previously described (*68*). Individual ZsGreen-1-NFAT reporter TCR transductants (20,000 cells/well) were cultured in the presence or absence of K562 cells expressing each Mel1 donor-derived HLA allele (50,000 cells/well) overnight, followed by evaluation of ZsGreen1 expression by flow cytometry. Cultures with 5 µg/ml of anti-mouse CD3ε antibody (clone 125-2C11, BD) were included in the assay as s positive control.

### Evaluation of responses to primary human islets and the PPI:15-24 peptide

Human islets (3,000–5,000 IEQs) from donors carrying HLA-A2 were obtained from the IIDP and cultured in 14 ml of CMRL medium (Gibco) supplemented with 10% FBS, 1× GlutaMax (Gibco), 1× penicillin-streptomycin (Sigma-Aldrich), 1 mM sodium pyruvate (Gibco), 10 mM HEPES (Gibco), and 0.25 μg/ml amphotericin B (Corning) in 10-cm Petri dishes at 37°C for 5 days before use in TCR transductant stimulation assays. Interferon-gamma (PeproTech) was added at 10 ng/ml for the final 2 days of culture. Islets were collected, washed with PBS, and dissociated into single cells by treatment with 0.05% trypsin-EDTA (Gibco) at 37°C for 10–15 minutes. Three to four TCR transductants engineered with a ZsGreen1 NFAT reporter (20,000 cells/well per clonotype) were pooled and cultured with or without islet cells (100,000 cells/well) in 200 μl of TCR transductant media in round-bottom 96-well plates overnight, followed by measurement of ZsGreen1 expression by flow cytometry. Anti-mouse CD3ε antibody (5 µg/ml; clone 125-2C11, BD) was included as a positive control.

To evaluate responses to the PPI:15–24 peptide (ALWGPDPAAA), individual TCR transductants (20,000 cells/well) were cultured with or without peptide (Genemed Synthesis) at concentrations ranging from 1 nM to 1 µM in the presence of K562 transductant cells expressing HLA-A*02:01 (50,000 cells/well) overnight, followed by measurement of ZsGreen1 expression by flow cytometry.

### Statistics

Data were analyzed using GraphPad Prism (GraphPad Software) and R (R Core Team). The specific statistical tests used are indicated in the corresponding text and figure legends. A P value <0.05 was considered statistically significant.

## Acknowledgments

We thank Dr. Teresa DiLorenzo for her insightful comments and valuable suggestions. We are thankful to organ donors and their families. This research was performed with the support of the Network for Pancreatic Organ donors with Diabetes (nPOD; RRID:SCR_014641), a collaborative T1D research project. The content and views expressed are the responsibility of the authors and do not necessarily reflect the official view of nPOD. Organ Procurement Organizations (OPO) partnering with nPOD to provide research resources are listed at http://www.jdrfnpod.org/for-partners/npod-partners/. Several human pancreatic islets were provided by the Alberta Diabetes Institute IsletCore at the University of Alberta in Edmonton (http://www.bcell.org/adi-isletcore.html) with the assistance of the Human Organ Procurement and Exchange (HOPE) program, Trillium Gift of Life Network (TGLN), and other Canadian organ procurement organizations. Islet isolation was approved by the Human Research Ethics Board at the University of Alberta (Pro00013094). All donors’ families gave informed consent for the use of pancreatic tissue in research. Another portion of human pancreatic islets were provided by the NIDDK-funded Integrated Islet Distribution Program (IIDP) (RRID:SCR _014387) at City of Hope, NIH Grant # U24DK098085.

## Author contributions

A.W.M., H.A.R., and M.N. conceived the study and designed the studies. A.M.A., L.G.L., J.M.B., A.H.S., R.C., A.M.L., M.D., and J.A.B.B. performed experiments. K.L.W. developed custom Python and R scripts. R.M. provided essential information on viral peptides. K.L.W., S.C.K., H.A.R., and M.N. analyzed the data. M.N. wrote the manuscript, and all authors reviewed and edited the manuscript.

---

**Supplementary Figure 1.**
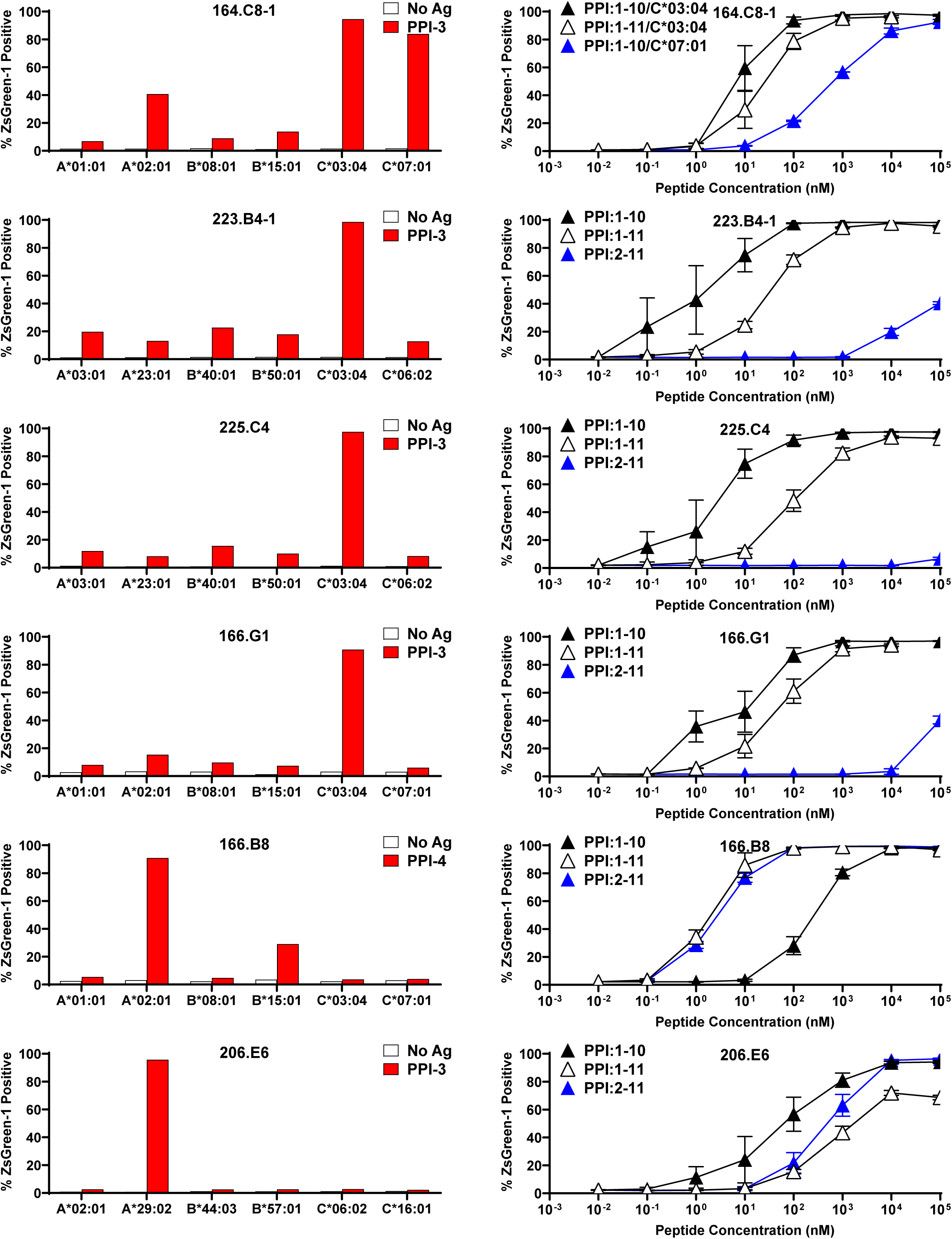
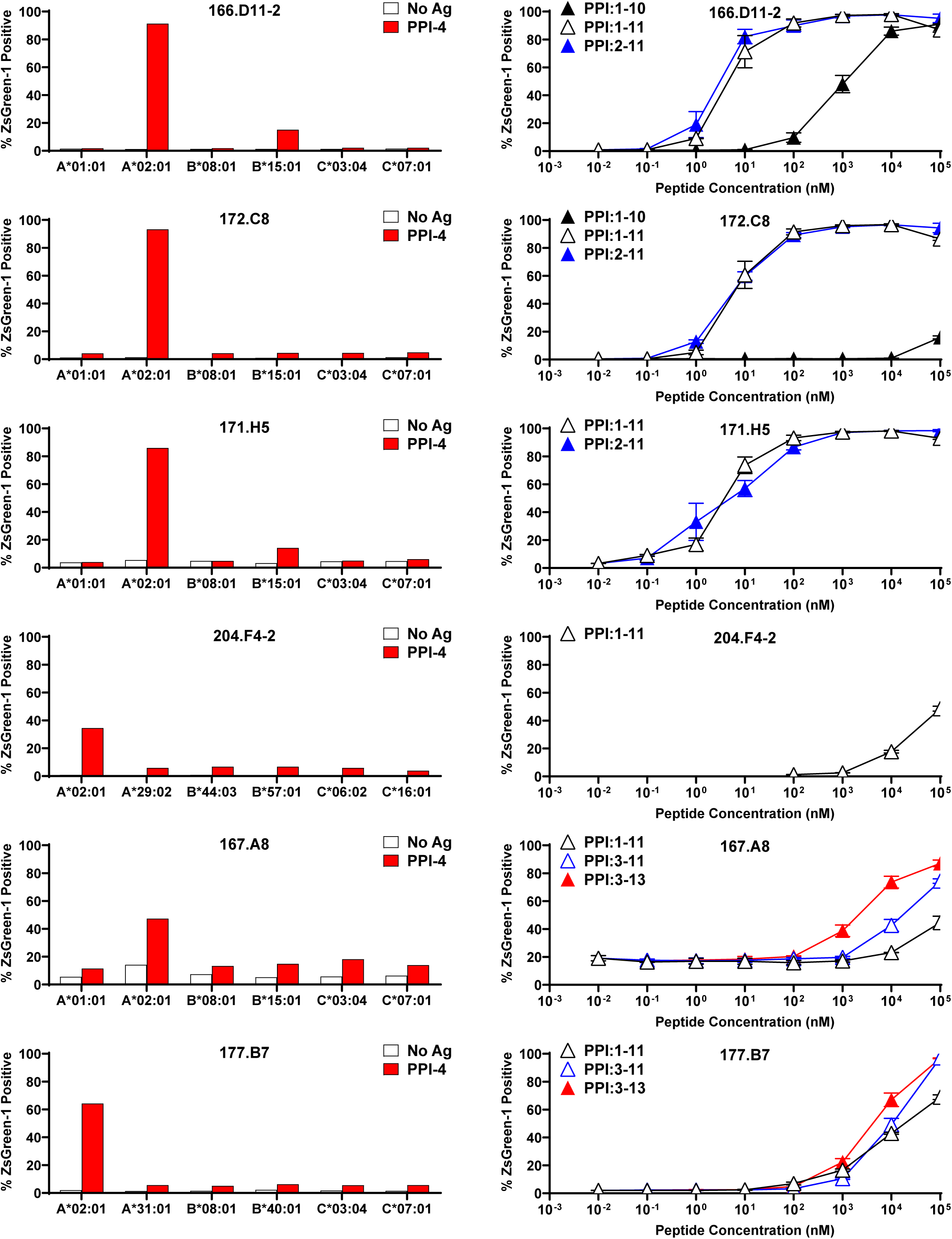
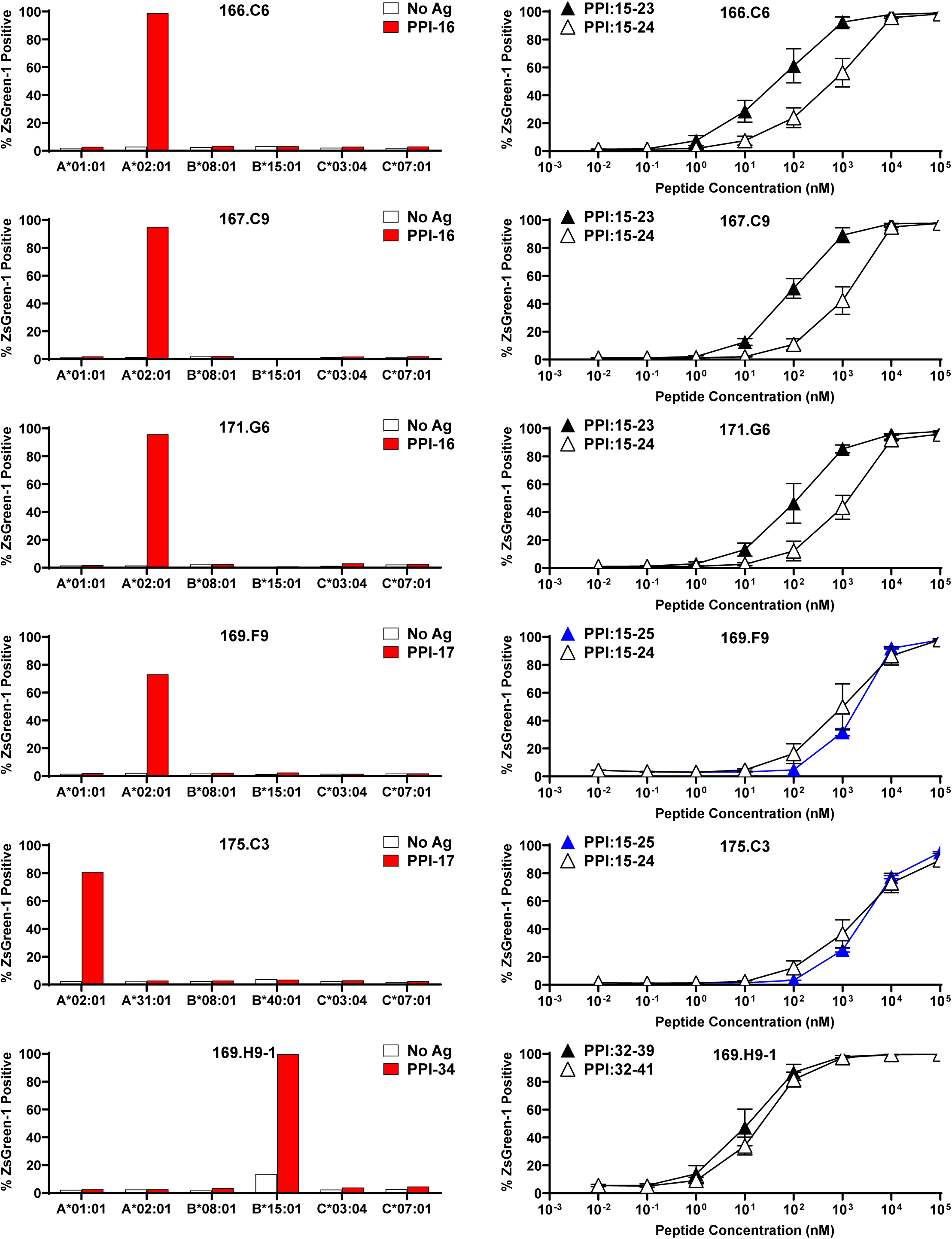
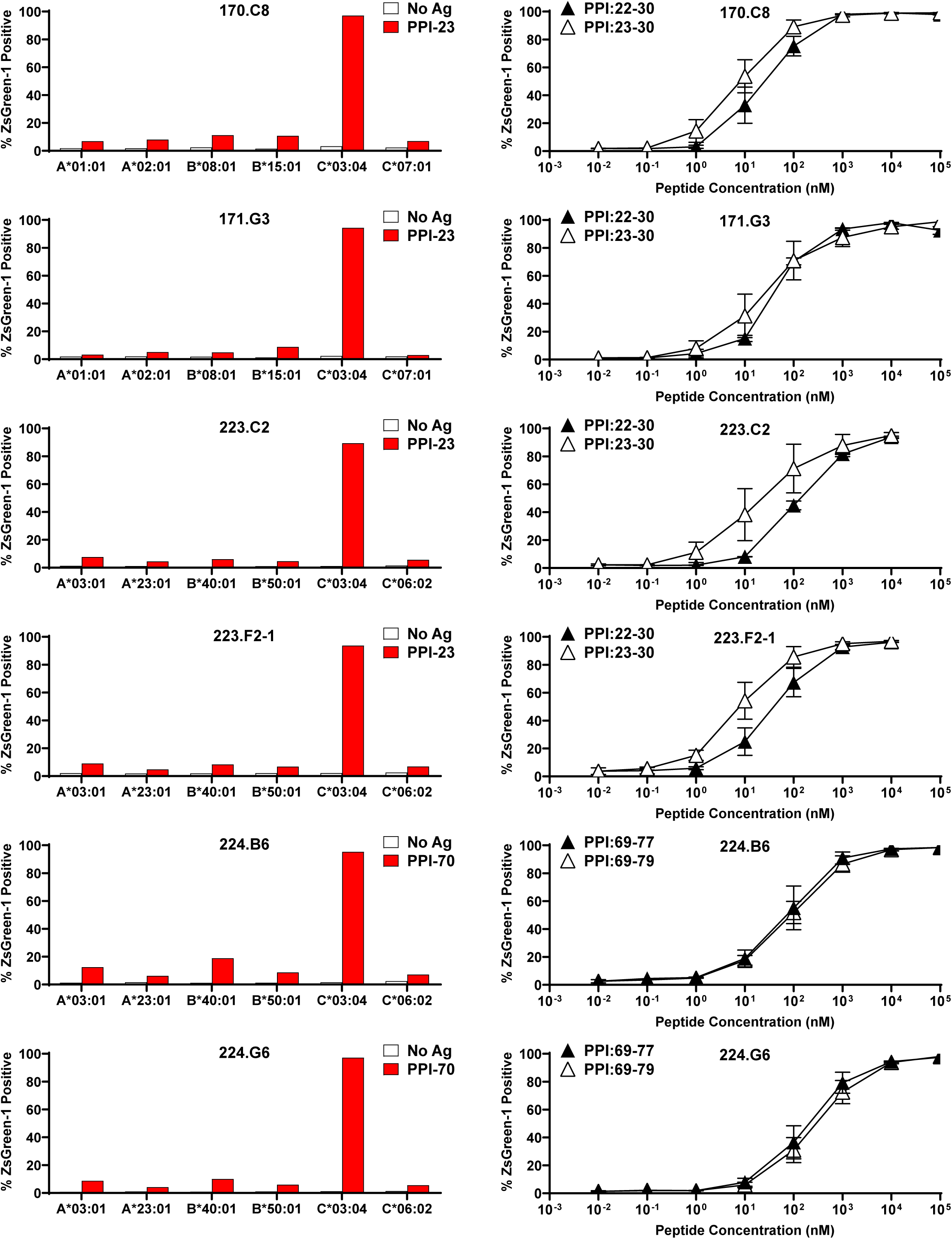
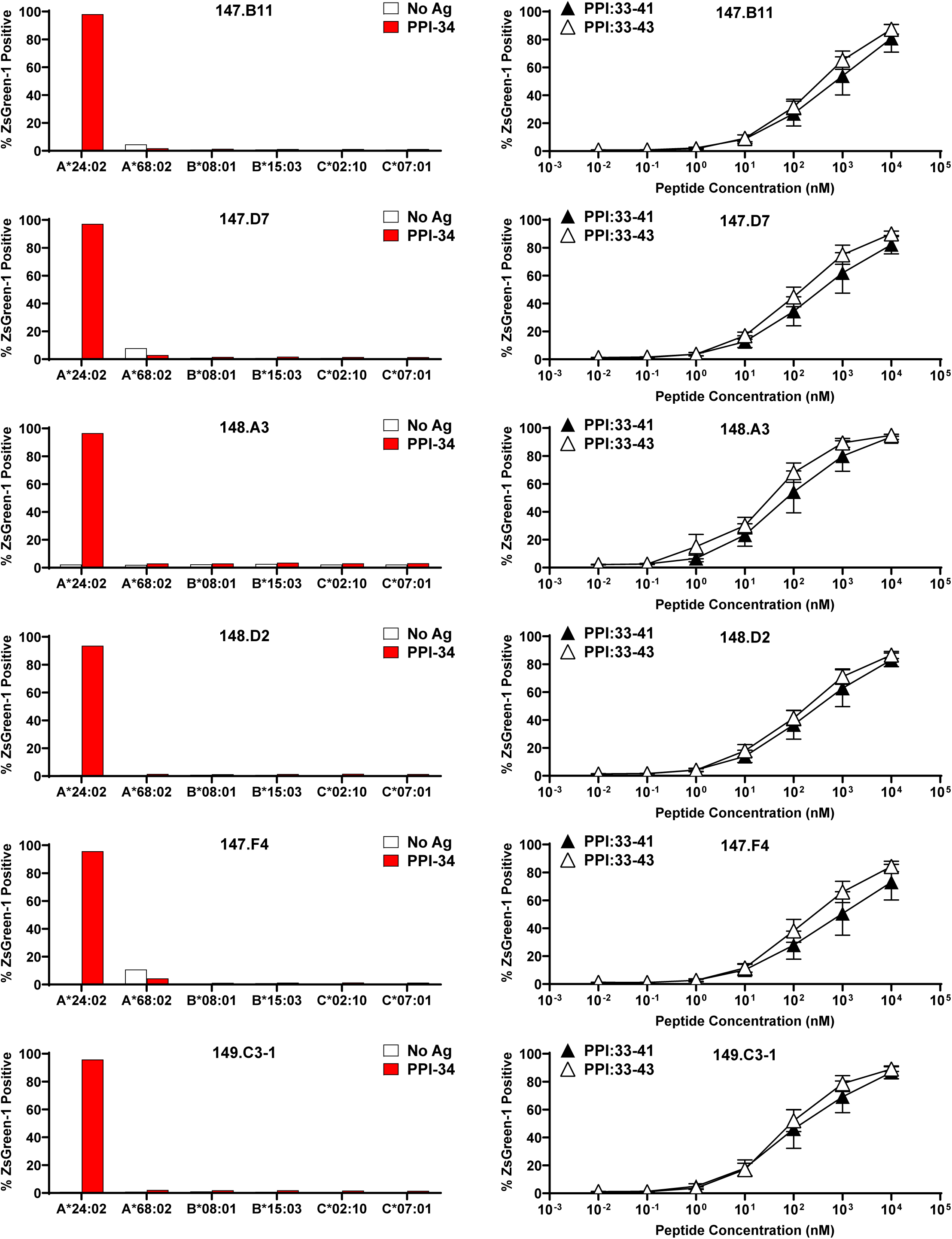
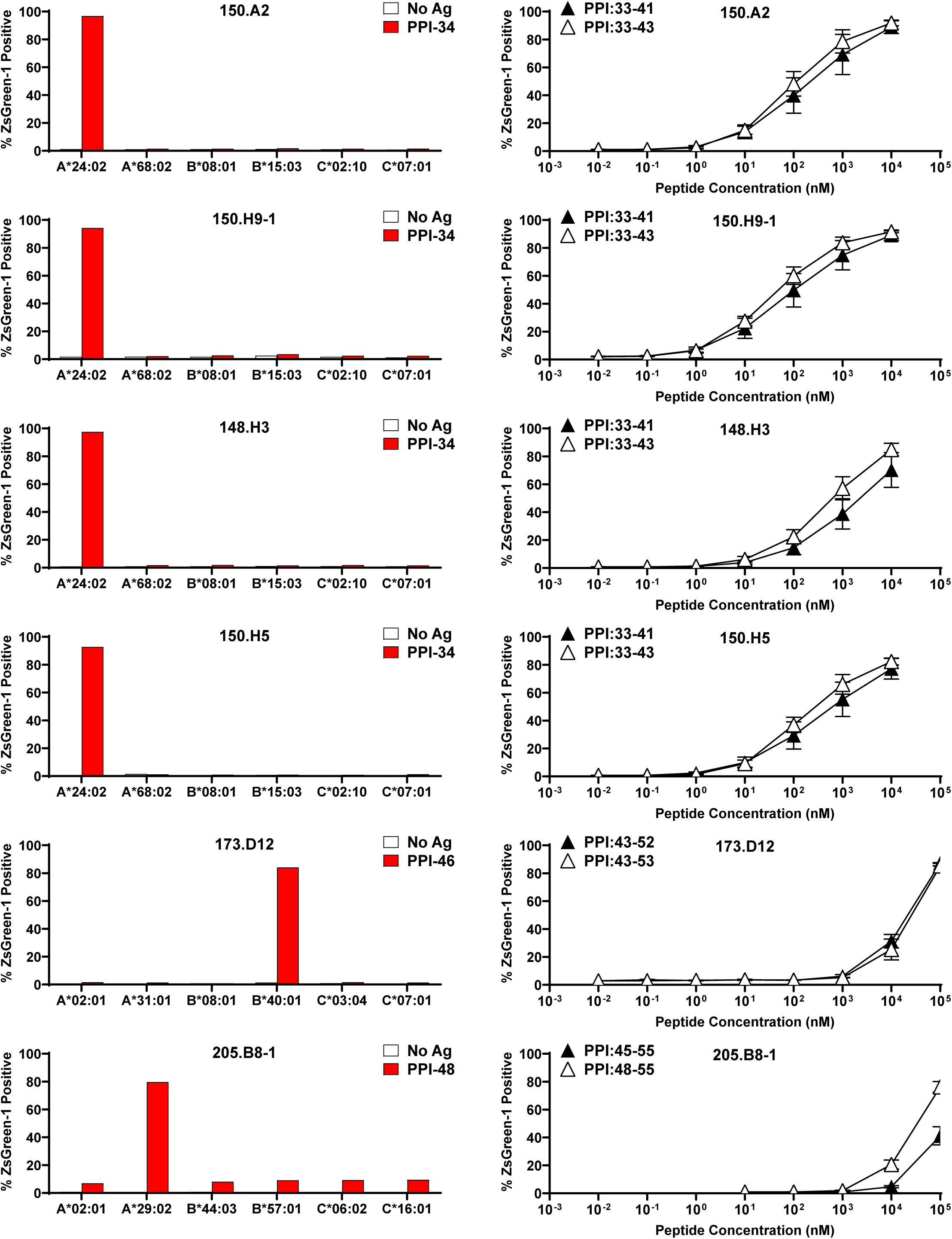
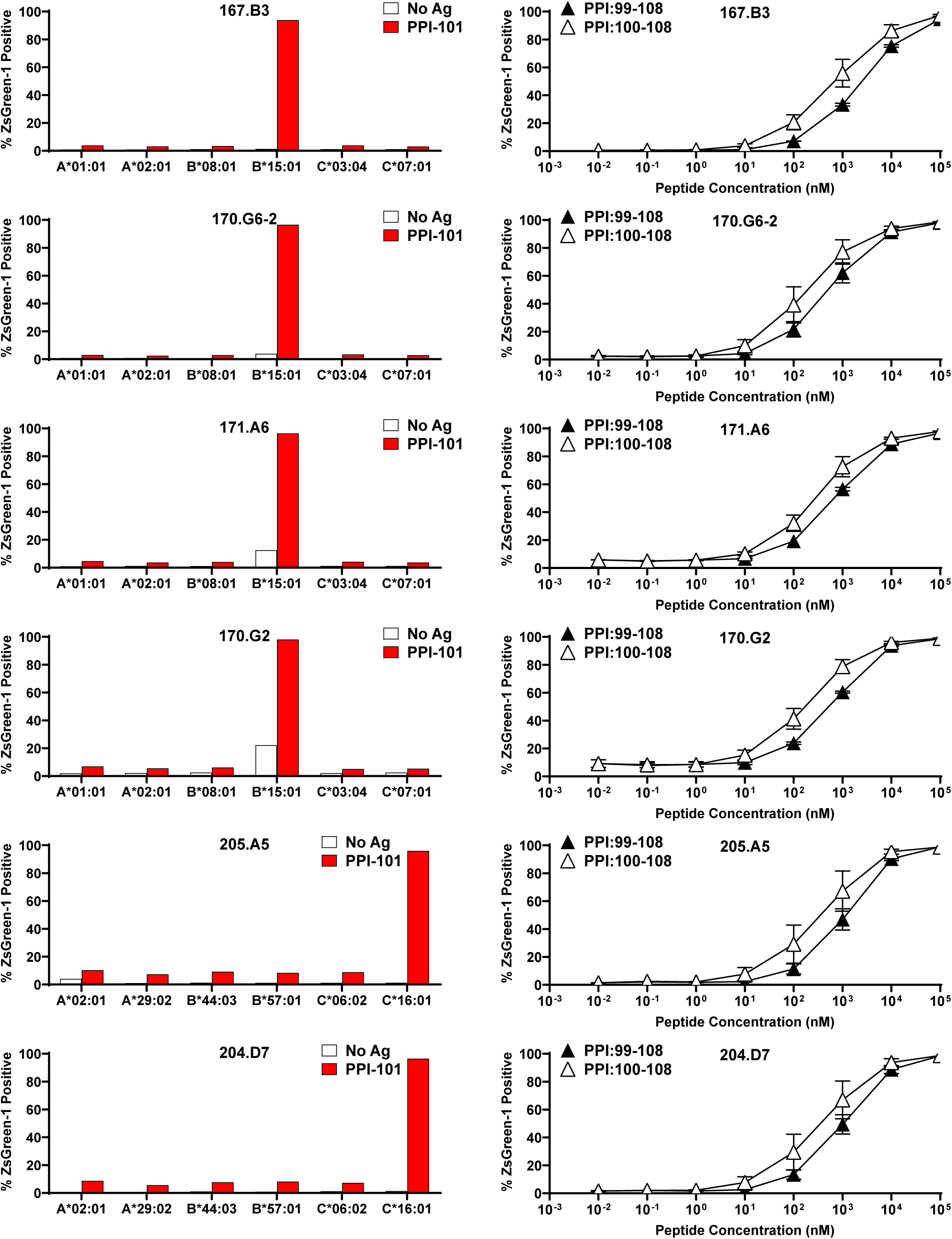
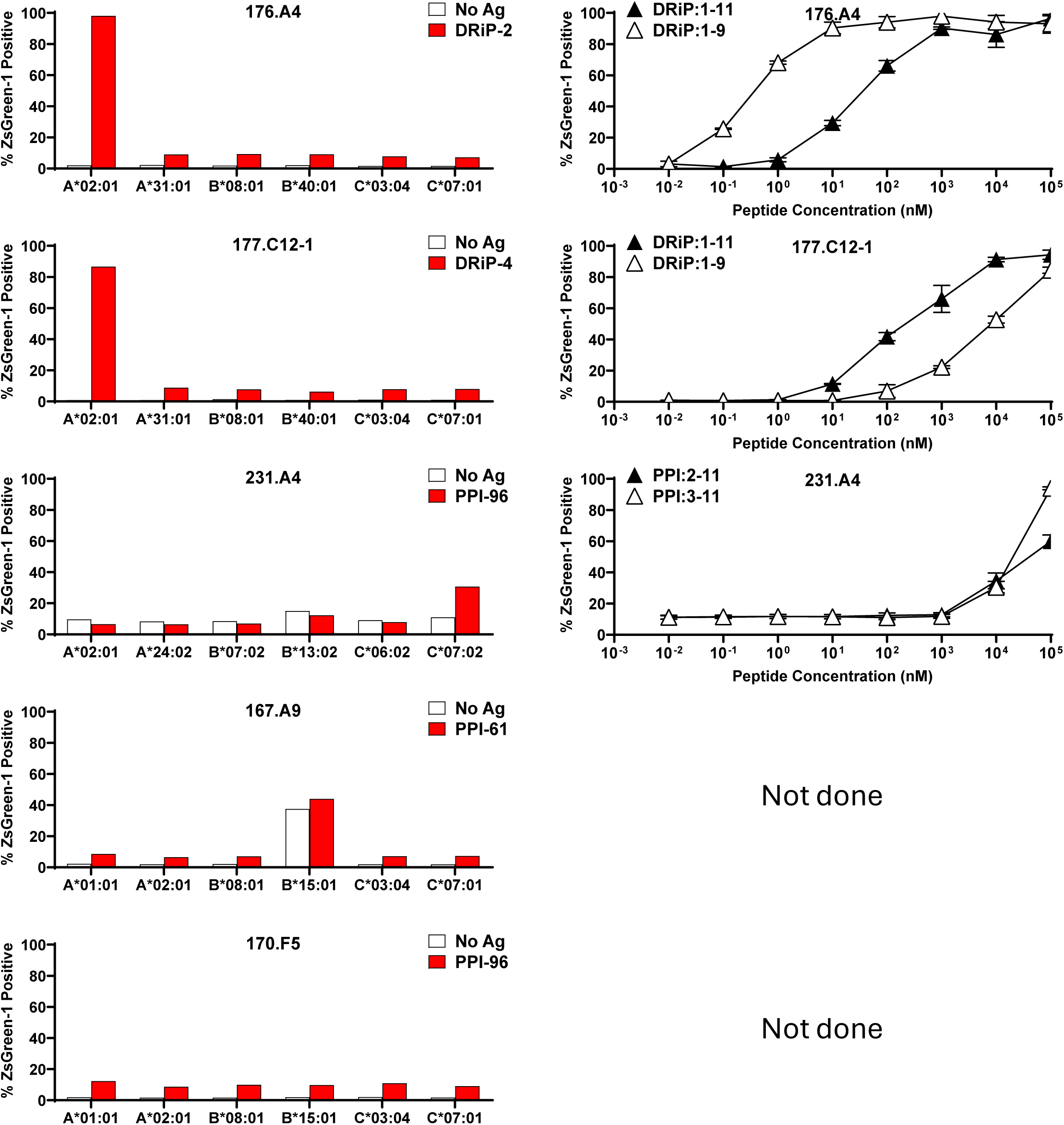
HLA restriction and titration analysis of PPI- and insulin-DRiP-reactive TCR transductants. **Left panels:** TCR transductants that responded to truncated peptide pools in the initial screen were tested using K562 cells expressing single donor-matched HLA class I molecules to confirm peptide specificity and define HLA restriction. Responses are shown as the percentage of ZsGreen1-positive cells following peptide stimulation. **Right panels:** For TCRs showing peptide-HLA-specific responses, TCR transductants were stimulated with cognate peptides from the corresponding pools across a range of concentrations in the presence of K562 cells expressing the defined HLA class I molecules. Activation was assessed by ZsGreen1 expression driven by an NFAT reporter, and mean values ± SEM from two or three independent experiments are shown.

**Supplementary Figure 2.**
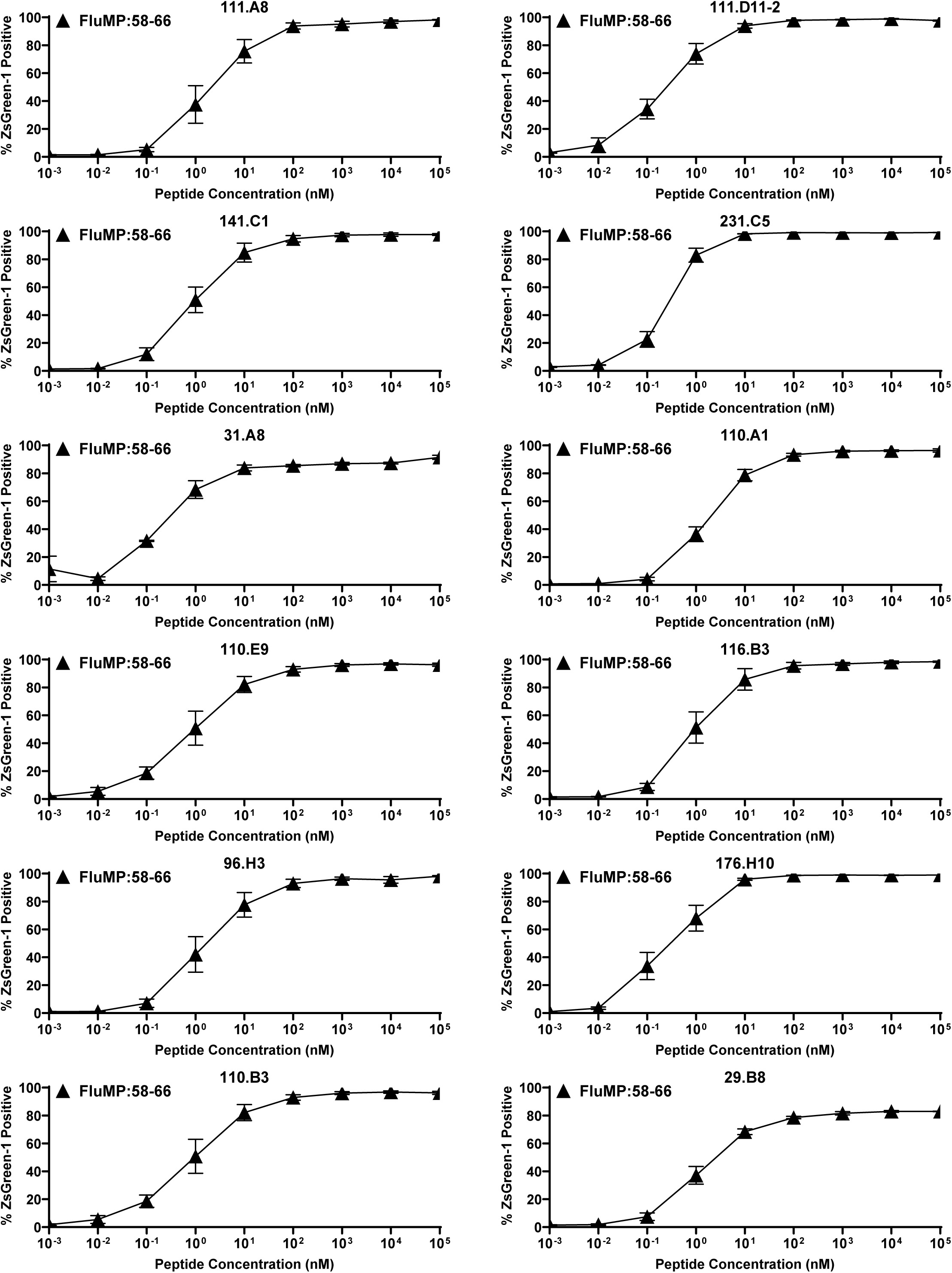
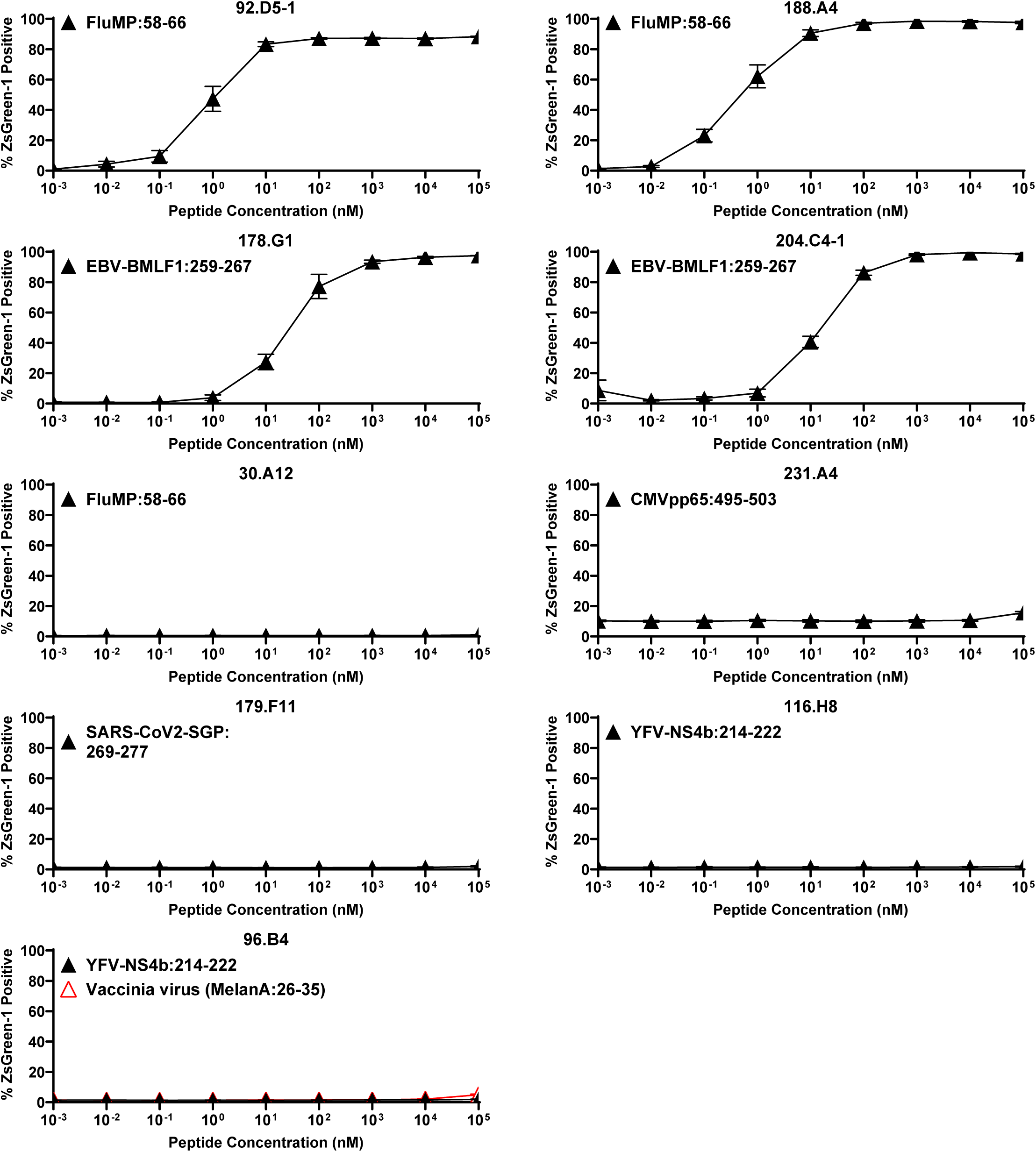
Titration analysis of transductants expressing public TCR clonotypes. Transductants expressing public TCR clonotypes identified from the pancreas were stimulated with cognate peptides reported in the Immune Epitope Database (IEDB) across a range of concentrations in the presence of K562 cells expressing the corresponding HLA class I molecules. Activation was assessed by ZsGreen1 expression driven by an NFAT reporter, and mean values ± SEM from three independent experiments are shown.

**Supplementary Figure 3.**
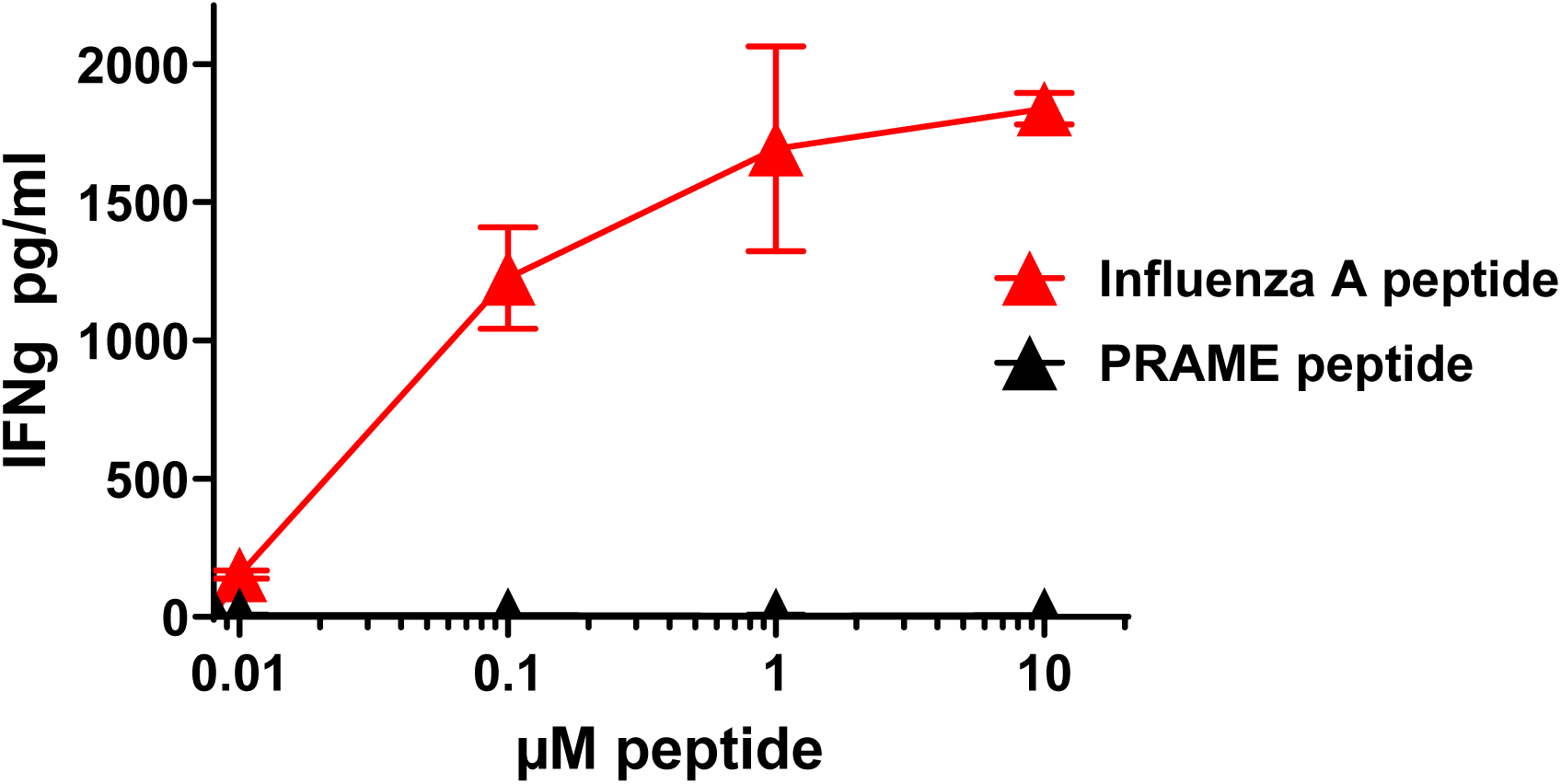
Influenza A peptide reactivity of islet-derived T cell clones from a T1D donor. Two T cell clones were established from T cell lines by 0.3 cells/well limiting dilution cloning derived from two individual islets hand-picked from enzyme-dispersed pancreas slice samples from an nPOD donor 6472 (10.25-year-old female with 4 years duration of T1D; RRIS:SAMN15879525). T cell lines that responded to the influenza A peptide were cloned by limiting dilution. Clones were cultured with the influenza A peptide (GILGFVFTL) or an islet-irrelevant control peptide, Preferentially Expressed Antigen of Melanoma (PRAME_425-433_ SLLQHLIGL), in the presence of HLA-A2-expressing C1R cells. Interferon-gamma secretion in culture supernatants was measured by ELISA. Experiments were performed in triplicate; representative results from one of the two clones are shown.

**Supplementary Figure 4.**
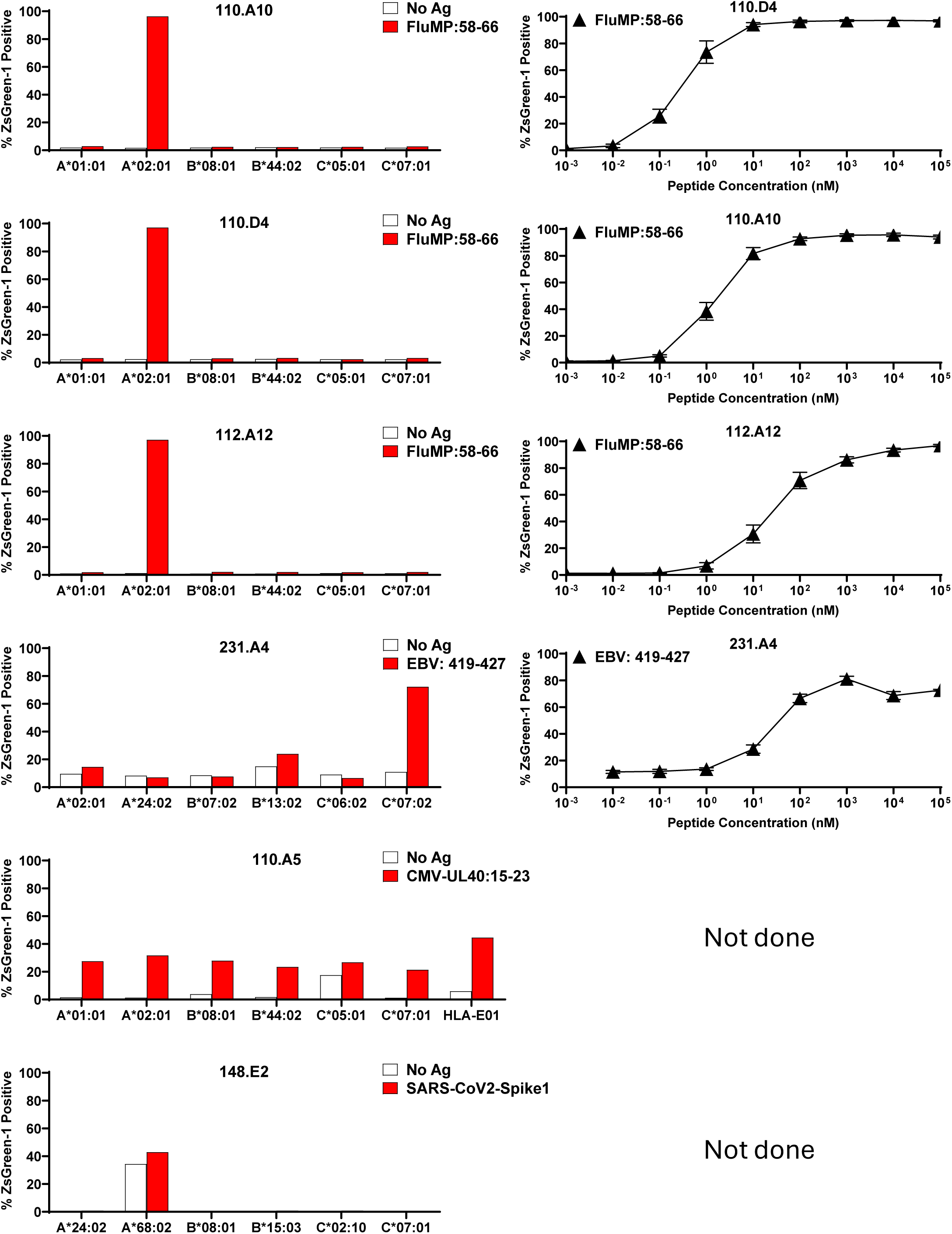
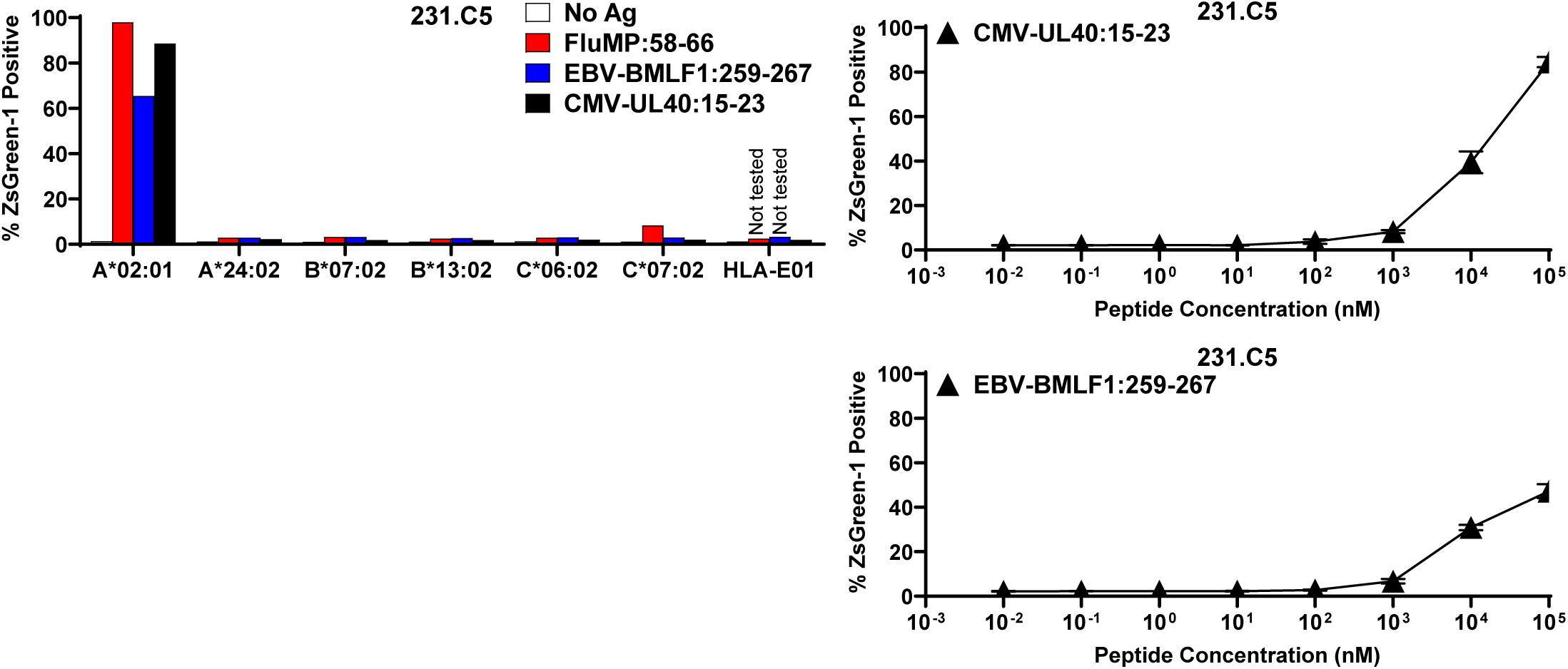
HLA restriction and titration analysis of viral peptide-reactive TCR transductants. **Left panels:** TCR transductants that responded to viral peptides in the initial screen were tested using K562 cells expressing single donor-matched HLA class I molecules to confirm peptide specificity and define HLA restriction. Responses are shown as the percentage of ZsGreen1-positive cells following peptide stimulation. **Right panels:** For TCRs showing peptide-HLA-specific responses, TCR transductants were stimulated with cognate peptides across a range of concentrations in the presence of K562 cells expressing the defined HLA class I molecules. Activation was assessed by ZsGreen1 expression driven by an NFAT reporter, and mean values ± SEM from three independent experiments are shown.

**Supplementary Figure 5.**
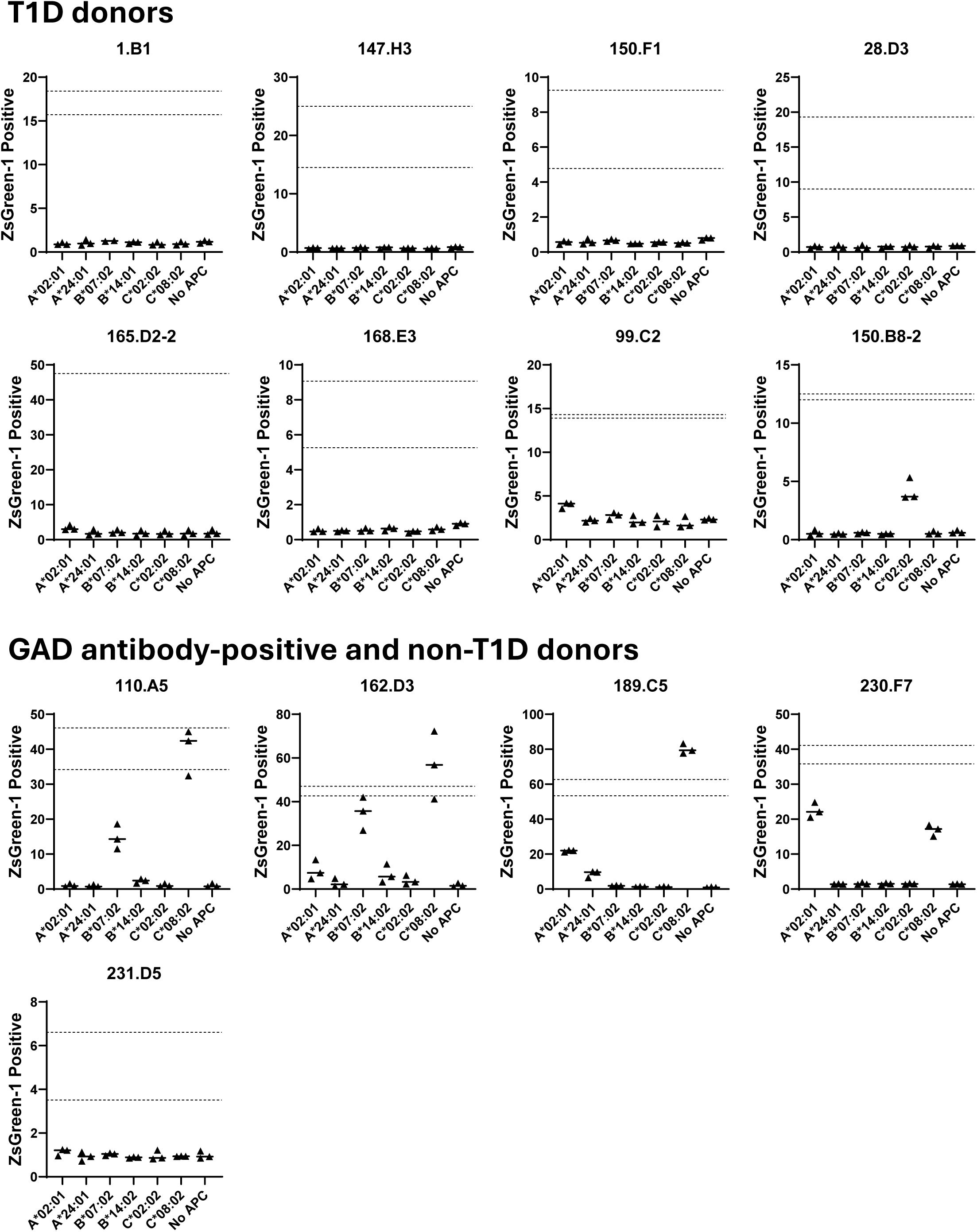
Testing allogeneic responses of sBC-reactive TCR transductants. TCR transductants defined as reactive to sBCs were tested for responses to K562 cells expressing individual HLA class I molecules carried by the sBC donor to assess potential alloreactivity. Activation was assessed by ZsGreen1 expression driven by an NFAT reporter, and mean values ± SEM from three independent experiments are shown.

## References

1. E. A. James, A. V. Joglekar, A. K. Linnemann, H. A. Russ, S. C. Kent, The beta cell-immune cell interface in type 1 diabetes (T1D). Molecular metabolism 78, 101809 (2023).

2. K. C. Herold et al., The immunology of type 1 diabetes. Nature reviews. Immunology 24, 435–451 (2024).

3. A. J. Dwyer, Z. R. Shaheen, B. T. Fife, Antigen-specific T cell responses in autoimmune diabetes. Frontiers in immunology 15, 1440045 (2024).

4. T. Delong, M. Nakayama, Epitope Hierarchy in Type 1 Diabetes Pathogenesis. Cold Spring Harb Perspect Med 15, (2025).

5. R. Mallone, C. Halliez, J. Rui, K. C. Herold, The β-Cell in Type 1 Diabetes Pathogenesis: A Victim of Circumstances or an Instigator of Tragic Events? Diabetes 71, 1603–1610 (2022).

6. D. L. Eizirik, F. Szymczak, R. Mallone, Why does the immune system destroy pancreatic β-cells but not α-cells in type 1 diabetes? Nature reviews. Endocrinology 19, 425–434 (2023).

7. E. A. James, R. Mallone, S. C. Kent, T. P. DiLorenzo, T-Cell Epitopes and Neo-epitopes in Type 1 Diabetes: A Comprehensive Update and Reappraisal. Diabetes 69, 1311–1335 (2020).

8. N. Amdare, A. W. Purcell, T. P. DiLorenzo, Noncontiguous T cell epitopes in autoimmune diabetes: From mice to men and back again. The Journal of biological chemistry 297, 100827 (2021).

9. J. Groegler, A. Callebaut, E. A. James, T. Delong, The insulin secretory granule is a hotspot for autoantigen formation in type 1 diabetes. Diabetologia, (2024).

10. S. S. Rich, H. Erlich, P. Concannon, in Diabetes in America, C. C. Cowie et al., Eds. (National Institute of Diabetes and Digestive and Kidney Diseases (US), Bethesda (MD), 2018).

11. J. A. Noble, Fifty years of HLA-associated type 1 diabetes risk: history, current knowledge, and future directions. Frontiers in immunology 15, 1457213 (2024).

12. K. Lipponen et al., Effect of HLA class I and class II alleles on progression from autoantibody positivity to overt type 1 diabetes in children with risk-associated class II genotypes. Diabetes 59, 3253–3256 (2010).

13. K. T. Coppieters et al., Demonstration of islet-autoreactive CD8 T cells in insulitic lesions from recent onset and long-term type 1 diabetes patients. The Journal of experimental medicine 209, 51–60 (2012).

14. J. A. Babon et al., Analysis of self-antigen specificity of islet-infiltrating T cells from human donors with type 1 diabetes. Nature medicine 22, 1482–1487 (2016).

15. S. Culina et al., Islet-reactive CD8(+) T cell frequencies in the pancreas, but not in blood, distinguish type 1 diabetic patients from healthy donors. Science immunology 3, (2018).

16. T. Rodriguez-Calvo, L. Krogvold, N. Amirian, K. Dahl-Jørgensen, M. von Herrath, One in Ten CD8(+) Cells in the Pancreas of Living Individuals With Recent-Onset Type 1 Diabetes Recognizes the Preproinsulin Epitope PPI(15-24). Diabetes 70, 752–758 (2021).

17. A. M. Anderson et al., Human islet T cells are highly reactive to preproinsulin in type 1 diabetes. Proceedings of the National Academy of Sciences of the United States of America 118, (2021).

18. T. Rodriguez-Calvo, S. J. Richardson, A. Pugliese, Pancreas Pathology During the Natural History of Type 1 Diabetes. Current diabetes reports 18, 124 (2018).

19. S. J. Richardson et al., Islet cell hyperexpression of HLA class I antigens: a defining feature in type 1 diabetes. Diabetologia 59, 2448–2458 (2016).

20. A. E. Wiedeman et al., Autoreactive CD8+ T cell exhaustion distinguishes subjects with slow type 1 diabetes progression. The Journal of clinical investigation 130, 480–490 (2020).

21. K. C. Herold et al., An Anti-CD3 Antibody, Teplizumab, in Relatives at Risk for Type 1 Diabetes. The New England journal of medicine, (2019).

22. R. Mallone et al., CD8+ T-cell responses identify beta-cell autoimmunity in human type 1 diabetes. Diabetes 56, 613–621 (2007).

23. A. Skowera et al., CTLs are targeted to kill beta cells in patients with type 1 diabetes through recognition of a glucose-regulated preproinsulin epitope. The Journal of clinical investigation 118, 3390–3402 (2008).

24. M. Scotto et al., Zinc transporter (ZnT)8(186-194) is an immunodominant CD8+ T cell epitope in HLA-A2+ type 1 diabetic patients. Diabetologia 55, 2026–2031 (2012).

25. N. Amdare et al., Islet-derived T cells from both mice and humans recognize conserved insulin A-chain peptides presented by HLA-C*03:04. Journal of immunology (Baltimore, Md. : 1950) 215, (2026).

26. M. P. Nekoua, E. K. Alidjinou, D. Hober, Persistent coxsackievirus B infection and pathogenesis of type 1 diabetes mellitus. Nature reviews. Endocrinology 18, 503–516 (2022).

27. A. Carré, F. Vecchio, M. Flodström-Tullberg, S. You, R. Mallone, Coxsackievirus and Type 1 Diabetes: Diabetogenic Mechanisms and Implications for Prevention. Endocr Rev 44, 737–751 (2023).

28. S. J. Richardson et al., Joint analysis of the nPOD-Virus Group data: the association of enterovirus with type 1 diabetes is supported by multiple markers of infection in pancreas tissue. Diabetologia 68, 1226–1241 (2025).

29. S. J. Micallef et al., INS(GFP/w) human embryonic stem cells facilitate isolation of in vitro derived insulin-producing cells. Diabetologia 55, 694–706 (2012).

30. S. Gonzalez-Duque et al., Conventional and Neo-antigenic Peptides Presented by β Cells Are Targeted by Circulating Naïve CD8+ T Cells in Type 1 Diabetic and Healthy Donors. Cell metabolism 28, 946–960.e946 (2018).

31. A. Carré et al., Interferon-α promotes HLA-B-restricted presentation of conventional and alternative antigens in human pancreatic β-cells. Nature communications 16, 765 (2025).

32. P. P. Nanaware et al., The antigen presentation landscape of cytokine-stressed human pancreatic islets. Cell reports 44, 115927 (2025).

33. K. Walters et al., Proteogenomic Discovery of Novel Open Reading Frames With HLA Immune Presentation on Human β-Cells. Diabetes 74, 2322-2336 (2025).

34. J. Champagne et al., Adoptive T cell therapy targeting an inducible and broadly shared product of aberrant mRNA translation. Immunity 58, 247–262.e249 (2025).

35. F. Vecchio et al., Coxsackievirus infection induces direct pancreatic β cell killing but poor antiviral CD8(+) T cell responses. Science advances 10, eadl1122 (2024).

36. T. Rodriguez-Calvo et al., Means, Motive, and Opportunity: Do Non-Islet-Reactive Infiltrating T Cells Contribute to Autoimmunity in Type 1 Diabetes? Frontiers in immunology 12, 683091 (2021).

37. A. M. Magnuson et al., Population dynamics of islet-infiltrating cells in autoimmune diabetes. Proceedings of the National Academy of Sciences of the United States of America 112, 1511–1516 (2015).

38. G. Christoffersson, G. Chodaczek, S. S. Ratliff, K. Coppieters, M. G. von Herrath, Suppression of diabetes by accumulation of non-islet-specific CD8(+) effector T cells in pancreatic islets. Science immunology 3, (2018).

39. J. W. Yewdell, Confronting complexity: real-world immunodominance in antiviral CD8+ T cell responses. Immunity 25, 533–543 (2006).

40. E. Assarsson et al., Kinetic analysis of a complete poxvirus transcriptome reveals an immediate-early class of genes. Proceedings of the National Academy of Sciences of the United States of America 105, 2140–2145 (2008).

41. A. V. Joglekar et al., T cell antigen discovery via signaling and antigen-presenting bifunctional receptors. Nature methods 16, 191–198 (2019).

42. T. Kula, et al., T-Scan: A Genome-wide Method for the Systematic Discovery of T Cell Epitopes. Cell 178, 1016-1028.e1013 (2019).

43. C. S. Dobson et al., Antigen identification and high-throughput interaction mapping by reprogramming viral entry. Nature methods 19, 449–460 (2022).

44. P. M. Bruno et al., High-throughput, targeted MHC class I immunopeptidomics using a functional genetics screening platform. Nature biotechnology 41, 980–992 (2023).

45. L. Wooldridge et al., A single autoimmune T cell receptor recognizes more than a million different peptides. The Journal of biological chemistry 287, 1168–1177 (2012).

46. G. Dolton et al., HLA A*24:02-restricted T cell receptors cross-recognize bacterial and preproinsulin peptides in type 1 diabetes. The Journal of clinical investigation 134, (2024).

47. K. Cerosaletti et al., Single-Cell RNA Sequencing Reveals Expanded Clones of Islet Antigen-Reactive CD4(+) T Cells in Peripheral Blood of Subjects with Type 1 Diabetes. Journal of immunology (Baltimore, Md. : 1950) 199, 323–335 (2017).

48. D. Kronenberg et al., Circulating preproinsulin signal peptide-specific CD8 T cells restricted by the susceptibility molecule HLA-A24 are expanded at onset of type 1 diabetes and kill β-cells. Diabetes 61, 1752–1759 (2012).

49. A. Skowera et al., β-cell-specific CD8 T cell phenotype in type 1 diabetes reflects chronic autoantigen exposure. Diabetes 64, 916–925 (2015).

50. A. Lledó-Delgado et al., Teplizumab induces persistent changes in the antigen-specific repertoire in individuals at risk for type 1 diabetes. The Journal of clinical investigation 134, (2024).

51. J. E. Slansky, M. Nakayama, Peptide mimotopes alter T cell function in cancer and autoimmunity. Semin Immunol 47, 101395 (2020).

52. S. I. Mannering et al., The insulin A-chain epitope recognized by human T cells is posttranslationally modified. The Journal of experimental medicine 202, 1191–1197 (2005).

53. M. van Lummel et al., Posttranslational modification of HLA-DQ binding islet autoantigens in type 1 diabetes. Diabetes 63, 237–247 (2014).

54. T. Delong et al., Pathogenic CD4 T cells in type 1 diabetes recognize epitopes formed by peptide fusion. Science (New York, N.Y.) 351, 711–714 (2016).

55. M. Buitinga et al., Inflammation-Induced Citrullinated Glucose-Regulated Protein 78 Elicits Immune Responses in Human Type 1 Diabetes. Diabetes 67, 2337–2348 (2018).

56. R. L. Baker et al., Hybrid Insulin Peptides Are Autoantigens in Type 1 Diabetes. Diabetes 68, 1830–1840 (2019).

57. X. Wan et al., The MHC-II peptidome of pancreatic islets identifies key features of autoimmune peptides. Nature immunology 21, 455–463 (2020).

58. N. Srivastava et al., A microenvironment-driven HLA-II-associated insulin neoantigen elicits persistent memory T cell activation in diabetes. Nature immunology 27, 82–97 (2026).

59. P. Bhattacharjee, et al., Proinsulin C-peptide is a major source of HLA-DQ8 restricted hybrid insulin peptides recognized by human islet-infiltrating CD4(+) T cells. PNAS Nexus 3, pgae491 (2024).

60. M. L. Marre et al., Modifying Enzymes Are Elicited by ER Stress, Generating Epitopes That Are Selectively Recognized by CD4(+) T Cells in Patients With Type 1 Diabetes. Diabetes **67**, 1356-1368 (2018).

61. E. K. Sims, R. G. Mirmira, C. Evans-Molina, The role of beta-cell dysfunction in early type 1 diabetes. Current opinion in endocrinology, diabetes, and obesity 27, 215–224 (2020).

62. B. O. Roep, S. Thomaidou, R. van Tienhoven, A. Zaldumbide, Type 1 diabetes mellitus as a disease of the β-cell (do not blame the immune system?). Nature reviews. Endocrinology 17, 150–161 (2021).

63. W. Wu et al., The Impact of Pro-Inflammatory Cytokines on Alternative Splicing Patterns in Human Islets. Diabetes 71, 116–127 (2021).

64. A. Coomans de Brachène et al., Interferons are key cytokines acting on pancreatic islets in type 1 diabetes. Diabetologia 67, 908–927 (2024).

65. F. Engin, D. L. Eizirik, Beyond bystanders: How β cell stress shapes the autoimmune response in T1D. Science translational medicine 18, eaed3762 (2026).

66. J. S. Kaddis, A. Pugliese, M. A. Atkinson, A run on the biobank: what have we learned about type 1 diabetes from the nPOD tissue repository? Current opinion in endocrinology, diabetes, and obesity 22, 290–295 (2015).

67. S. E. Mann et al., Multiplex T Cell Stimulation Assay Utilizing a T Cell Activation Reporter-Based Detection System. Frontiers in immunology 11, 633 (2020).

68. L. G. Landry, S. E. Mann, A. M. Anderson, M. Nakayama, Multiplex T-cell Stimulation Assay Utilizing a T-cell Activation Reporter-based Detection System. Bio-protocol 11, e3883 (2021).

69. G. Tosato, J. I. Cohen, Generation of Epstein-Barr Virus (EBV)-immortalized B cell lines. Curr Protoc Immunol Chapter 7, 7.22.21–27.22.24 (2007).

70. H. A. Russ et al., Controlled induction of human pancreatic progenitors produces functional beta-like cells in vitro. The EMBO journal 34, 1759–1772 (2015).

71. M. E. Brown et al., Human stem cell-derived β cells expressing an optimized CD155 reduce cytotoxic immune cell function for application in type 1 diabetes. Science advances 11, eadx9755 (2025).

72. J. M. Barra et al., Cryopreservation of Stem Cell-Derived β-Like Cells Enriches for Insulin-Producing Cells With Improved Function. Diabetes 73, 1687–1696 (2024).

